# Altered empathy processing in frontotemporal dementia A task-based fMRI study

**DOI:** 10.1101/2024.03.21.586051

**Authors:** Olof Lindberg, Tie-Qiang Li, Cecilia Lind, Susanna Vestberg, Ove Almkvist, Mikael Stiernstedt, Anita Ericson, Nenad Bogdanovic, Oskar Hansson, Luke Harper, Eric Westman, Caroline Graff, Theofanis Tsevis, Peter Mannfolk, Håkan Fischer, Gustav Nilsonne, Predrag Petrovic, Lars Nyberg, Lars-Olof Wahlund, Alexander F Santillo, the Swedish FTD initiative

**Affiliations:** Department of Neurobiology, Care Sciences and Society, Karolinska Institute, Division of Clinical Geriatrics, Centre for Alzheimer Research, Neo, 14183 Huddinge, Sweden; Department of Clinical Science, Intervention, and Technology, Karolinska Institute, Sweden; Department of Medical Radiation and Nuclear Medicine, Karolinska University Hospital, Sweden; Department of community medicine and rehabilitation, geriatrics Umeå university, Umeå university, Sweden; Department of Psychology, Lund University, Lund, Sweden; Department of Psychology, Stockholm, Sweden; Umeå center for Functional Brain Imaging (UFBI), Umeå University, Sweden; Clinical Memory Research Unit, Department of Clinical Sciences Malmö, Lund University, Lund, Sweden; Memory Clinic, Skåne University Hospital, Lund, Sweden; Karolinska university hospital, Stockholm, Sweden; Department of Neurobiology, Care Sciences and Society, Division of Neurogeriatrics, Center for Alzheimer Research, Karolinska Institutet, Solna, Sweden; Department of Medical Imaging and Physiology, Skåne University Hospital, Lund, Sweden; Stockholm University Brain Imaging Centre (SUBIC), Stockholm, Sweden; Aging Research Center, Karolinska Institutet, Stockholm, Sweden; Center for Psychiatric Research, Department of Clinical Neuroscience, Karolinska Institute, Sweden; Center for Cognitive and Computational Psychiatry, Department of Clinical Neuroscience, Karolinska Institute, Sweden; Department of Radiation Sciences, Umeå University, Umeå, Sweden; Department of Medical and Translational Biology, Umeå University, Umeå, Sweden

**Keywords:** Frontotemporal dementia, Empathy, Functional MRI, Empathic concern

## Abstract

A lack of empathy, and particularly its affective components, is a core symptom of behavioural variant frontotemporal dementia (bvFTD). Visual exposure to images of a needle pricking a hand (pain condition) and Q-tips touching a hand (control condition) is an established functional magnetic resonance imaging (fMRI) paradigm used to investigate empathy for pain (EFP; pain condition minus control condition). EFP has been associated with increased blood oxygen level dependent (BOLD) signal in regions known to become atrophic in the early stages in bvFTD, including the anterior insula and the anterior cingulate. We therefore hypothesized that patients with bvFTD would display altered empathy processing in the EFP paradigm. Here we examined empathy processing using the EFP paradigm in 28 patients with bvFTD and 28 sex and age matched controls. Participants underwent structural MRI, task-based and resting-state fMRI. The Interpersonal Reactivity Index (IRI) was used as a measure of different facets of empathic function outside the scanner. The EFP paradigm was analysed at a whole brain level and using two regions-of-interest approaches, one based on a metanalysis of affective perceptual empathy versus cognitive evaluative empathy and one based on the control’s activation pattern. In controls, EFP was linked to an expected increase of BOLD signal that displayed an overlap with the pattern of atrophy in the bvFTD patients (insula and anterior cingulate). Additional regions with increased signal were the supramarginal gyrus and the occipital cortex. These latter regions were the only ones that displayed increased BOLD signal in bvFTD patients. BOLD signal increase under the affective perceptual empathy but not the cognitive evaluative empathy region of interest was significantly greater in controls than in bvFTD patients. The control’s rating on their empathic concern subscale of the IRI was significantly correlated with the BOLD signal in the EFP paradigm, as were an informant’s ratings of the patient’s empathic concern subscale. This correlation was not observed on other subscales of the IRI or when using the patient’s self-ratings. Finally, controls and patients showed different connectivity patterns in empathy related networks during resting-state fMRI, mainly in nodes overlapping the ventral attention network. Our results indicate that reduced neural activity in regions typically affected by pathology in bvFTD is associated with reduced empathy processing, and a predictor of patient’s capacity to experience affective empathy.

## Introduction

Behavioural variant frontotemporal dementia (bvFTD) is a syndrome of a neurodegenerative disease commonly affecting the anterior frontal, temporal and insular cortices.^1^ Symptoms of bvFTD include a combination of changes in behaviour (disinhibition, lack of motivation/ apathy, repetitive behaviour and altered eating habits), and cognitive dysfunction (mainly in social cognition, executive functions and language), concomitant to a lack of insight.^1,2^ Important drivers of the changes in behaviour are loss of social cognitive functions such as emotional recognition, understanding of social norms, mentalization (“theory of mind”), moral reasoning, and empathic function, which occur together to varying degrees (for reviews see, ^3,4^). Loss of empathy, and particularly its affective aspects, appear to be independent of decrease in the other socioemotional abilities and general cognition.^5–7^ Close relatives and caregivers often report a lack of warmth and connectedness in patients with bvFTD.^8^ From a diagnostic perspective, loss of empathy is an early symptom in bvFTD, with high specificity, allowing differentiation from other neurological diseases.^9,10^ Furthermore, a lack of empathy is one of the six recognised core clinical symptoms of the current International Consensus Criteria for the diagnosis of bvFTD.^2^ Clinically, loss of empathy is an independent predictor of functional loss and caregiver burden in bvFTD.^11,12^

Brain atrophy in bvFTD has been shown to initially occur in the anterior insula (AI) and the anterior cingulate cortex (ACC).^13,14^ Whilst there is heterogeneity in the pattern of atrophy and heterogeneity in respect to underlying neuropathology of bvFTD, studies have demonstrated a consistent core of neurodegeneration involving the AI and ACC across all cases of bvFTD.^15,16^ This characteristic pattern of neurodegeneration is further reflected in reduced functional connectivity of the AI and ACC, which may be identified using resting state functional MRI ^17,18^. The AI and the ACC (its posterior part, sometimes referred to as midcingulate cortex, MCC) have been shown to be central nodes in the ventral attention network (VAN),^19^ also referred to as the “salience network”.^14,20^ The VAN is one of several intrinsically connected resting-state networks.^21^ The VAN is proposed to be involved in the detection and integration of emotional and sensory stimuli within motivational, social, and cognitive contexts. These convergent processes culminate in the highest level of integration, forming what has been referred to as a ’global emotional moment’ in the fronto-insular area, particularly within the right hemisphere.^20,22^ Dysfunction of the VAN, primarily caused by atrophy and reduced neural activity of the AI and ACC, has been demonstrated in bvFTD, and has been tied to specific clinical characteristics such as disinhibition, apathy,^17,20,23,24^ and disruption of a long range of socioemotional functions, including empathy.^25–27^

Empathy is a multidimensional construct that is supported by several interacting brain networks.^28^ Conceptually it is commonly divided into two domains, an affective domain (affective empathy), defined as the vicarious experience of other’s sensorimotor or emotional state and a cognitive domain (cognitive empathy), largely synonymous with “theory of mind” or “mentalizing”, defined as the ability to understand the perspective of another person while maintaining a distinction between one’s own and the other person’s mental state.^28,29^ A prototypical instrument that aims to capture both affective empathy and cognitive empathy is the Interpersonal Reactivity Index (IRI).^30^ The IRI is a questionnaire where empathy is rated along four subscales/subcomponents,^31^ of which the “empathic concern” (EC) and the “personal distress” (PD) subscales are thought to relate to affective empathy. These traits are associated with specific brain circuits that are of fundamental importance for the ability to experience different components of affective empathy.^31,32^ The other two subscales, the “perspective taking” (PT) and “fantasy subscale” (FS) relate to the mentalizing ability and therefore represents cognitive empathy. In bvFTD only affective empathy has been observed to be independent of other cognitive domains, while cognitive empathy has been found to depend on other abilities, particularly executive function.^5–8^

The neuroanatomical correlates of affective and cognitive empathy have been extensively investigated during the last decades using functional magnetic resonance imaging (fMRI). Recently a meta-analysis included 103 task-based paradigms that assessed cognitive empathy and 85 paradigms that assessed affective empathy,^33^ found that affective empathy was associated with increased blood-oxygen-level-dependent (BOLD) signal in areas such as bilateral insula, supramarginal gyrus, supplementary motor area and midcingulate cortex, predominantly involving regions belonging to the VAN.^33^ Additionally reduced affective empathy has been observed in patients with lesions in areas belonging to the VAN, such as the AI.^34,35^ One study found that lesions to the AI, but not to the ACC impaired patients in empathic pain perception (one aspect of affective empathy),^36^ suggesting that the AI has a decisive role for the ability to experience empathy for pain (EFP). Cognitive empathy, on the other hand, have in fMRI studies been associated with activity in midline areas of the prefrontal cortex, precuneus and parietal regions, predominantly overlapping with the default mode network (DMN).^19^

Affective empathy can in turn be subdivided into distinctive empathy processes disentangled using different experimental paradigms. The possibly most studied is the EFP paradigm (induced by watching other individuals experiencing pain), which is associated with increase activation in insula bilaterally and the caudal ACC.^37–40^ This paradigm suggests partially overlapping information processing for experiencing subjective pain and for watching other individuals in pain. Although this overlap has been shown to not represent the same activation matrix^41^, abundant evidence suggests that this network is involved in affective aspects of empathy.^33^ An example of such overlap is the insula that has an important role in EFP, and an equally important role in processing of nociception intensity and localisation.^42,43,44^

Thus, previous studies have revealed that the prototypical pattern of atrophy bvFTD involve regions that are active during an affective perceptual empathic response. However, to our knowledge, brain activation in an empathic response has not been linked to the core neuroanatomical signatures of bvFTD. Here, we aimed to:

- measure dynamic brain activity while empathizing in individuals with bvFTD using a well-established task-based fMRI paradigm.^45–47^

- explore the relationship of the activation pattern in the fMRI task with empathic function outside the scanner, using the IRI.

- compare patients with controls regarding the strength of resting-state connectivity between regions involved in a normal empathy-associated activation.

- analyze the diagnostic potential of the task-based fMRI assessment of empathy compared to that of cortical atrophy.

## Materials and methods

### Participants

Individuals were recruited between 2015-2022 at three Memory Clinics in Sweden: Skåne University Hospital Malmö, Norrlands University Hospital Umeå and Karolinska University Hospital in Huddinge. Inclusion criteria were a diagnosis of bvFTD according to International Behavioral Variant FTD Consortium Criteria^2^ (either possible, probable or definite bvFTD) following multidisciplinary assessment including clinical examination, caregiver interview, clinical neuropsychological examination, neuroimaging and lumbar puncture. Cerebrospinal fluid (CSF) was analysed for amyloid β, tau and phosphorylated tau (p-tau). Due to the cognitive demands of the task-based paradigm only individuals with mild dementia as specified by a clinical dementia rating (CDR) ≤ 1,^48^ or a mini mental status examination (MMSE) ≥ 21 were included.^49^ Additional inclusion criteria included the availability of a reliable informant, and proficiency in the Swedish language was required of both the patients and informants. As the EFP activation pattern overlaps with that of subjective pain,^39^ and some patients with bvFTD may have altered sensation to pain (hypo- or hyperesthesia,^50^), this data was recorded from most participants and their informants. Participants with a combination of CSF amyloid β (either amyloid β 42 or amyloid β 42/40 ratio) below and CSF p-tau above local laboratory reference values, indicative of Alzheimer’s disease, were excluded from study. Genetic screening for bvFTD associated genes was not done consecutively but where clinically indicated.

In addition to neuropsychological testing performed during clinical assessment of included patients, did a majority patients and controls perform a set of tests as a part of the present study. Global cognition was measured using the MMSE,^49^ executive function/working memory was assessed with the Digit Span backward tests, attention/processing speed was assessed with the Digit Span forward (both from WAIS-IV,^51^) and psychomotor speed was assessed using the Trail Making Test part A.^52^ A convenience sampled control group of healthy individuals matched according to age and sex with the patient group were included. They underwent the same study examinations as the patients. All neuropsychological data were standardized to *z*-scores. Mean score of controls were set to 0. A flowchart of the enrolled individuals is provided in the Supplementary Fig. 1.

The study was approved by the Regional Ethics Review board in Stockholm 2013- 04-19 (diary number 2013279-31). Written consent in accordance with the Helsinki declaration was obtained from all participants included in the study.

### The interpersonal reactivity index

The interpersonal reactivity index (IRI) is a 28-items questionary that aims to measure a persońs ability to react to the observed experience of another person with four subscales: One) Perspective taking (PT), the tendency to spontaneously adopt the psychological point of view of others, two) Fantasy (FS), the respondent’s tendencies to transpose themselves imaginatively into the feelings and actions of fictitious characters in books, movies or plays, three) Empathic concern (EC), the “other-oriented” feelings of personal unease and anxiety for others in distress, and four) Personal Distress (PD), that taps in on feelings of personal anxiety and unease in tense interpersonal settings.^30^ Rating is performed along a Likert scale with five levels. One characteristic of patients with bvFTD is loss of insight.^2^ Therefore, in addition to patients rating themselves on the IRI an informant (in most cases a close relative living with the patient or someone with regular and frequent interaction with the patient) rated the patients, as customary in research on bvFTD using the IRI’.^7,27,53^ Controls rated themselves. Pearsońs correlation was used to study the relationship between ratings on the IRI subscale and mean BOLD signal during EFP.

### The fMRI task

During the fMRI paradigm participants were exposed to images of a needle pricking a hand (pain condition) or a Q-tip touching a hand (control condition)^54^. Both conditions were displayed in 20 trials in a semi-randomized order (Fig.1; for Swedish translation of text used in the paradigm see Supplementary material).

**Figure 1.**
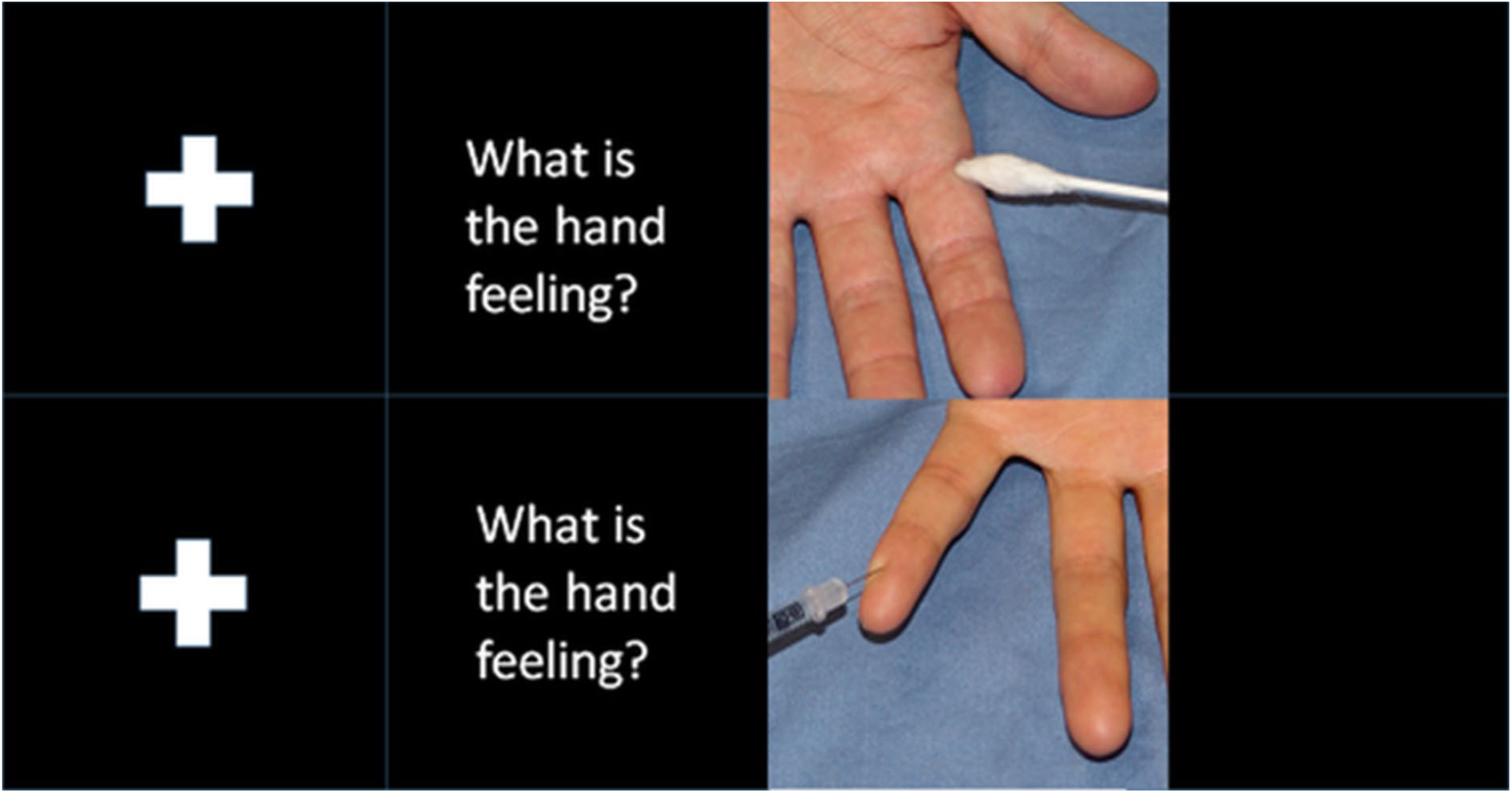
The experimental fMRI paradigm. The figure shows the images displayed during the fMRI-experiment using the empathy of pain paradigm. First was a fixation cross displayed for 2-4 seconds, then the text “What is the hand feeling?” for 3 seconds, and then an image displaying either a hand touched by a Q-tip or penetrated by a needle for 3.5 seconds, then a new sequence is initiated with a fixation cross.

### MRI Image acquisition

Structural and functional MRI data was acquired with a 64-channel head coil in two 3T Prisma MRI scanners (Siemens MAGNETOM Prisma, Siemens Healthcare, Erlangen, Germany) in Lund and in Stockholm, and one Discovery MR750 (General Electric system, GE Healthcare, United States) using a 32-channel head coil in Umeå. In the Prisma scanners task-based fMRI images were acquired with a TR=3000 ms TE=34 ms, FA 90 degrees, with an in-plane resolution of 2.3mm*2.3mm and slice thickness of 2.3 mm. In the GE MRI scanner task-based fMRI images were acquired with a TR=3000 ms TE=34 ms, FA=85 degrees, with an in-plane resolution of 2.38*2.38 and slice thickness of 2.3 mm. Parameters of the structural 3-D T1- weighted images, resting sting-state sequence and image quality control procedures are detailed in the Supplementary material.

### Image analysis

The analysis of regional brain volume and cortical thickness was performed using FreeSurfer 6 (https://surfer.nmr.mgh.harvard.edu). This analysis was performed for two reasons: 1) To evaluate whether the patients included in this study displayed the expected pattern of cortical thinning, and 2) to evaluate whether decreased cortical thickness was associated with decreased BOLD signal during EFP. For method description and the analysis of structural data see Supplementary material.

### Task-based fMRI

Analysis of task-based fMRI data was performed using FEAT v6.00 (FSL, FMRIB, Oxford, UK). Raw fMRI scans were pre-processed before statistical analysis using the following steps: motion correction using MCFLIRT,^55^ removal of nonbrain tissue using BET,^56^ spatial smoothing using a full width at half maximum of 5mm gaussian kernel. Time series statistical analysis and improved linear model (FILM) with local autocorrection was also performed ^57^. Finally, images were registered to the structural image and then to standard MNI space using FLIRT affine registration with 12 degrees of freedom. Relative displacement (head motion between consequentially acquired volumes) and absolute displacement (head motion in relation to the reference volume) was reported in FEAT.

### Whole brain analysis

Group-level fMRI analysis was performed using FMRIB Local Analysis of Mixed Effects (FLAME 1&2).^58,59^ Group level *z* (gaussianized *t*) statistical images were performed using a threshold of *z*>3.1, *p*<0.05 (whole-brain cluster-wise-corrected data). At individual level, *z* (gaussianized *t*) statistical images were produced. Values of *z*>2.3, *p*<0.05 (whole-brain cluster-wise-corrected data) was considered significant. The EFP contrast (pain condition minus control condition) was evaluated.

#### Regions of interest (ROI) analysis

The regions of interest-based analysis involved two main approaches. First we defined two sets of ROIs based on a previous neuroimaging meta-analysis that included 50 fMRI studies on empathy, separating affective perceptual empathy from cognitive evaluative empathy.^40^ One ROI was defined for each of these empathy conditions. Affective perceptual empathy was most commonly associated with increased BOLD signal in the right anterior insula extending to the right inferior frontal gyrus, the left insula and, and the right supplementary motor area (SMA). Cognitive evaluative empathy was associated with increased BOLD signal in the left anterior and midcingulate cortex and the left insula (Table 2 in Fan et al.^40^). We extracted mean BOLD signal using 5mm-spheres at MNI coordinates in which peak activation in the meta-analysis was identified in cognitive evaluative empathy (two yellow spheres in Supplementary in Fig. 2) and affective perceptual empathy (three red spheres in Supplementary Fig. 2). Mean BOLD signal under the spheres included in affective perceptual empathy (the affective perceptual ROI) and cognitive evaluative empathy (the cognitive evaluative empathy ROI) was compared between patients and controls.

**Figure 2.**
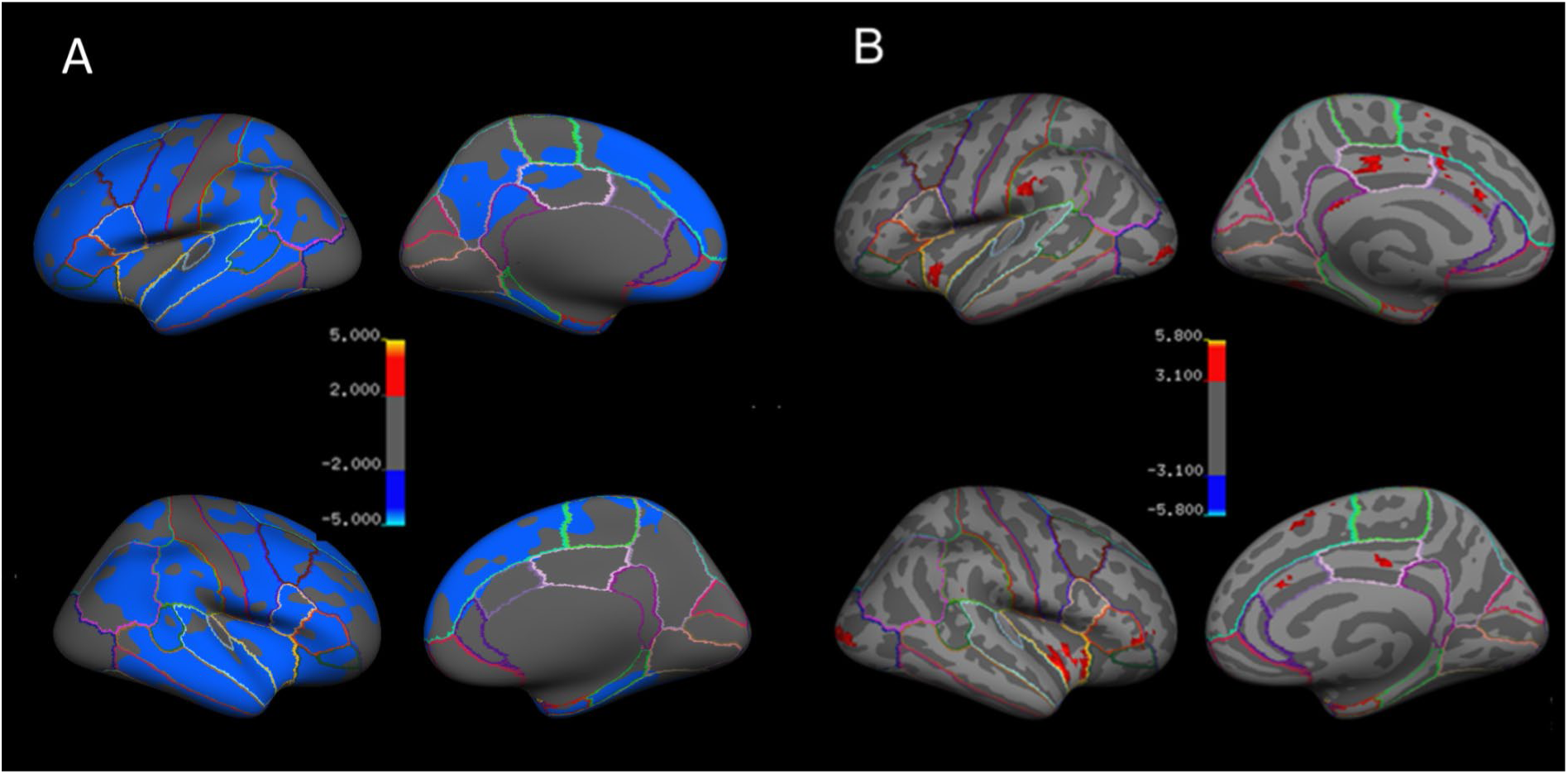
Cortical thickness relative to controls, and significant BOLD-signal in controls. Results displayed on an inflated cortical surface on fsaverage in FreeSurfer. Coloured lines denote the border of different cortical regions as defined in Desikan-Killany atlas.^76^ **A)** Reduced cortical thickness relative to controls in behavioural variant of frontotemporal dementia indicated by blue colour. Colour bar denotes -log10 *p-values*. Dark blue = *p<*0.01; light blue = *p=*0.00001. **B)** Regions with significantly increased BOLD signal during the empathy of pain contrast in the control group. Red colour indicates significant BOLD signal. The colour bar denotes *z-scores* red colour *z*>3.1, yellow *z*>5.

Second, we created one control activation ROI (CA-ROI) that encompassed 12 5mm spheres centred at the peak activation of the 12 areas with significant BOLD signal during the EFP contrast in the controls (Supplementary Fig. 3). This ROI was used to test empathic function in the patient group, as it mirrors the expected normal activation pattern in our experiment. The Featquery tool (https://fsl.fmrib.ox.ac.uk/fsl) was used to extract mean percent signal change in the EFP contrast across all participants. The ROI-based analyses were evaluated using general linear models (GLM) in SPSS (Armonk, NY: IBM Corp). Graphs were plotted using in Statistica (TIBCO Software Inc. version 13. http://tibco.co). In the total sample no covariates were included as the patient and control group was matched for relevant factors such as age and sex. In the sensitivity analysis relevant covariates were included and reported for each specific analysis below.

#### Sensitivity analyses

MRI scanner was not a significant predictor of BOLD signal under any of the investigated ROIs or of mean left or right cortyical thickness in patients investigated by different sites. MRI scanner was thus not included in the general linear models. We, however, evaluated the robustness of our results in several sensitivity analyses, in which we investigated the potential confounding effects of 1) Movement in the scanner 2) difference in cortical thickness between patients and controls, and 3) MRI scanners (Supplementary sensitivity analysis one to five).

### Resting-state fMRI connectivity analyses

Resting state intrinsic connectivity of brain regions that were activated during EFP in controls were assessed using Pearson’s correlation coefficients. Correlation between the average time courses of the 12 5 mm spheres that together constitute the CA-ROI was calculated. Differences between bvFTD patients and controls were evaluated using a two-sample t-test on these matrices. The Bonferroni approach was employed for multiple comparison correction. In order to explore the intrinsic connectivity network of each of the 12 task-defined CA-ROIs, Pearson’s correlation coefficients maps were computed between each ROI’s average time course and every other resting-state-fMRI data voxel, comparing the strength of correlation between patients and controls. Methods description of these analyses is detailed in the Supplementary material.

## Results

Patients with bvFTD (n=28) and control subjects (n=28) were included in the study (Table 1). All definite bvFTD cases were either carriers of pathogenic mutations in the chromosome 9 open reading frame 72 (*C9orf72*) gene, or the microtubule-associated protein tau (*MAPT*) gene, in the progranulin (*GRN*) gene. According to consensus criteria^2^, nine patients fulfilled criteria for definite bvFTD, 18 for probable and one for possible bvFTD. Two patients declined to undergo CSF analysis. Not all bvFTD did manage to perform the IRI self-rating, or the IRI self-rating was judged to be of insufficient quality (for example due to perseverations). Image quality control and the number of participants that completed IRI and neuropsychological testing are detailed in the flowchart (Supplementary Fig.1).

**Table 1.** Demographic, interpersonal reactivity index and neuropsychological test data.

|  | **CTL** | **bvFTD** |
| --- | --- | --- |
| Men/Women | 12/16 | 13/15 |
| Number in each MRI scanner | LU 13, KI 15 | LU 21, KI 2, UME 5 |
| Age | 67.59(7.44) | 66.74(6.74) |
| MMSE | 29.7(0.58) | 24.71(4.97)** |
| **IRI-ratings** | *Mean(SD)* | *Mean(SD)* |
| FS self-rating | 2.99(0.71) | 2.74(0.82) |
| EC self-rating | 3.11(0.28) | 3.28(0.75) |
| PT self-rating | 3.50(0.37) | 3.44(0.71) |
| PD self-rating | 3.05(0.42) | 3.03(0.97) |
| FS informants rating |  | 2.47(0.56) |
| EC informants rating |  | 3.14(0.35) |
| PT informants rating |  | 2.75(0.36) |
| PD informants rating |  | 2.90(0.51) |
| **Neuropsychological test** | *Mean z-score (SD)* | *Mean z-score (SD)* |
| Digit span forward | 0(1) | -0.41(0.87) |
| Digit-span backward | 0(1) | -0.58(1.01) |
| Trail making A | 0(1) | -2.63(4.06)** |

CTL= controls; bvFTD = behavioural variant Frontotemporal Dementia; *n* = number; Age = Age at scan; MMSE = mini mental state examination; *SD* = standard deviation; IRI = interpersonal reactivity index; IRI subscales: FS = fantasy; EC = empathic concern; PT = perspective taking; PD = personal distress. LU = Lund; KI = Karolinska; UME = Umeå; \*\* = $p < 0.01$ , \* = $p < 0.05$ in Student-T test.

Patients performed significantly worse than controls on Trail Making A test (controls mean*=*0.0 *SD=*1.0 bvFTD *mean=* -2.9*, SD=* 3.8*, p*<0.01) assessing visual attention and cognitive flexibility, but not on other neuropsychological tests (Table 1). Altered pain sensation was reported in two individuals with bvFTD. They displayed reduced BOLD signal under the CA-ROI (*F*[1. 16]=5.39, *p*=0.03), but not in the two meta-analysis ROIs compared patients without (or not reported) altered pain sensation.

### Cortical thickness and subcortical volume analyses

Patients displayed reduced cortical thickness in areas of the frontal, temporal, parietal and insular lobes compared to control subjects (Fig. 2A, Supplementary Table 1). Reduced thickness was observed in several regions that showed empathy related activation in controls, as depicted in Fig. 2B. Further to this, patients displayed reduced volume in several subcortical nuclei, detailed in Supplementary Table 2.

### Task-based fMRI

#### Whole-brain analysis

The control subjects displayed significantly increased BOLD signal in the EFP contrast (the pain condition minus control condition) in 12 areas, depicted in Fig. 3A and listed in Table 2. Regions of significantly increased BOLD signal overlapped the VAN in 7 locations. These included regions; Nr.1. Right posterior orbitofrontal cortex/AI; Nr. 3. Left AI; Nr.4. left dorsal ACC; Nr.5. left supramarginal gyrus; Nr.6. left supplementary motor area; Nr.8. right supramarginal gyrus; Nr.11. right pars opercularis in the inferior frontal gyrus. One cluster was located in the DMN (Nr.2. bilateral posterior cingulate), whilst three regions were located in the visual network (Nr.7. right occipital pole, Nr.9. left occipital pole & Nr.10. Occipital fusiform gyrus) and one region was located in the frontoparietal network (Nr.12. right frontal pole) as defined in Yeo et al 2011^19^ (Supplementary Fig. 3 & Table 2). In the patient group, no significant activations were observed in regions normally activated in the EFP contrast such as insula and ACC. The patients displayed significantly increased BOLD signal in the left occipital cortex and the left supramarginal gyrus, overlapping the visual network and VAN, respectively (Fig. 3B & Table 2.). If the two patients with altered pain perception were excluded, one additional cluster with significant BOLD signal during EFP was observed in patients, located in the posterior cingulate gyrus (data not shown). No significant activation differences between patients and controls were observed during EFP in the whole-brain analysis.

**Figure 3.**
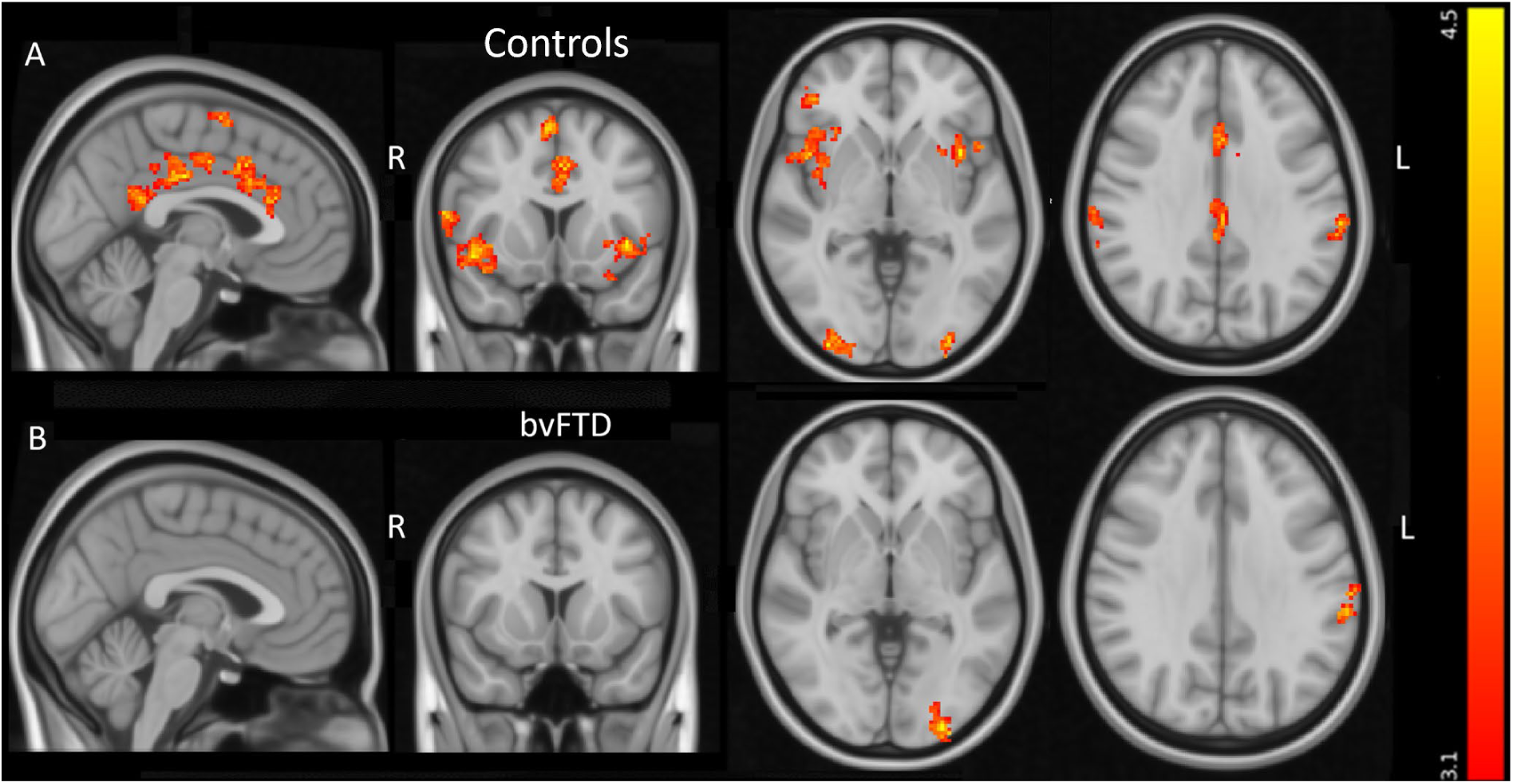
Areas with significant BOLD signal in the empathy of pain contrast. Areas with significant increased BOLD signal in control subjects (**A**) and in behavioural variant frontotemporal dementia (**B**) in the empathy of pain contrast. The colour bar displays z-scores (red *z*>3.1, yellow *z*=4.5). L = left; R = right. From left to right sagittal, coronal and axial slices at MNI coordinates *x* = -2, *y* =12, *z* = -2. Last axial slice at MNI coordinates *x* = 62, *y* = -32, *z* =32.

**Table 2.** Areas displaying activation during the empathy of pain in controls and in patients with bvFTD.

| Controls | Size | *p* | *z*-max | x | y | z | Area |
| --- | --- | --- | --- | --- | --- | --- | --- |
| 1 | 817 | 2.7e-11 | 5.58 | 42 | 22 | -6 | 49% Frontal orbital, 14% frontal operculum, 12% insula |
| 2 | 503 | 5.96e-08 | 5.19 | 0 | -24 | 34 | 96% Cingulate gyrus (posterior) |
| 3 | 435 | 3.58e-07 | 4.65 | -38 | 12 | -2 | 81% Insular cortex, |
| 4 | 372 | 1.97e-06 | 4.34 | -2 | 12 | 40 | 50% Cingulate / paracingulate cortex, 39% anterior |
| 5 | 270 | 4.51e-05 | 4.58 | -64 | -30 | 30 | 72% Supramarginal, 10% postcentral |
| 6 | 219 | 0.000249 | 4.49 | -4 | 0 | 62 | 84% Supplementary Motor Area |
| 7 | 155 | 0.0026 | 4.81 | 34 | -94 | -2 | 61% Occipital pole |
| 8 | 142 | 0.00435 | 4.05 | 68 | -24 | 30 | 30% Supramarginal gyrus |
| 9 | 135 | 0.00577 | 4.77 | -32 | -92 | -4 | 26% Occipital pole lateral |
| 10 | 107 | 0.0188 | 4.07 | -32 | -60 | -12 | 13% Occipital fusiform |
| 11 | 94 | 0.0335 | 4.27 | 62 | 12 | 14 | 16% Pars opercularis |
| 12 | 94 | 0.0335 | 4.07 | 42 | 40 | -2 | 44% Frontal pole, 4% Pars opercularis |
| bvFTD | Size | *P* | *z*-max | x | y | z | Area |
| 1 | 143 | 0.00418 | 5.47 | -26 | -96 | -4 | 52% Occipital pole, 6% Lateral occipital |
| 2 | 116 | 0.0127 | 4.2 | -62 | -32 | 32 | 73% Supramarginal Anterior division |

Controls = control subjects; bvFTD = patients with the behavioural variant of frontotemporal dementia; Size = number of voxels in cluster; *p* = value of cluster; x, y, z = MNI coordinates; z-stats = max z-value in cluster; Areas = percentage of anatomical areas contained in cluster.

### ROI-based analyses

#### The affective perceptual empathy and the cognitive evaluative empathy ROIs

Controls displayed increased activity in the affective perceptual empathy ROI in the EFP contrast (pain *mean* = 51.02 [*SD*=46.02] vs. no pain *mean* = 30.16 [*SD*=46.38], *t* = -3.88, *p*<0.01) but not in the cognitive evaluative empathy ROI (pain *mean* = 48.35 [*SD*=44.65] vs. no pain *mean*=41.73 [SD=49.95], *t*=-1.59, *p*=0.14). Patients displayed no increased activity in the affective perceptual empathy ROI (pain *mean*=14.19 [*SD*=45.08] vs. no pain *mean* =12.80 [*SD*=47.13], *t* = -0.44, *p*=0.66) or in the cognitive evaluative empathy ROI (pain *mean* =14.19 [*SD*=48.83] vs. no pain *mean* =11.87 [*SD* = 52.26], *t* = 0.38, *p* = 0.71). In a comparison between the groups, patients displayed decreased BOLD signal in the EFP contrast under the affective perceptual empathy ROI (*F*[1. 54] = 9.2, *p*<0.01; *η*2 = 0.15) compared with the controls, as depicted in Fig. 4**A**. No difference was identified between the groups in the cognitive evaluative empathy ROI, (*F*[1. 54]=2.50, *p*=0.12,) as depicted in Fig. 4**B**.

**Figure 4.**
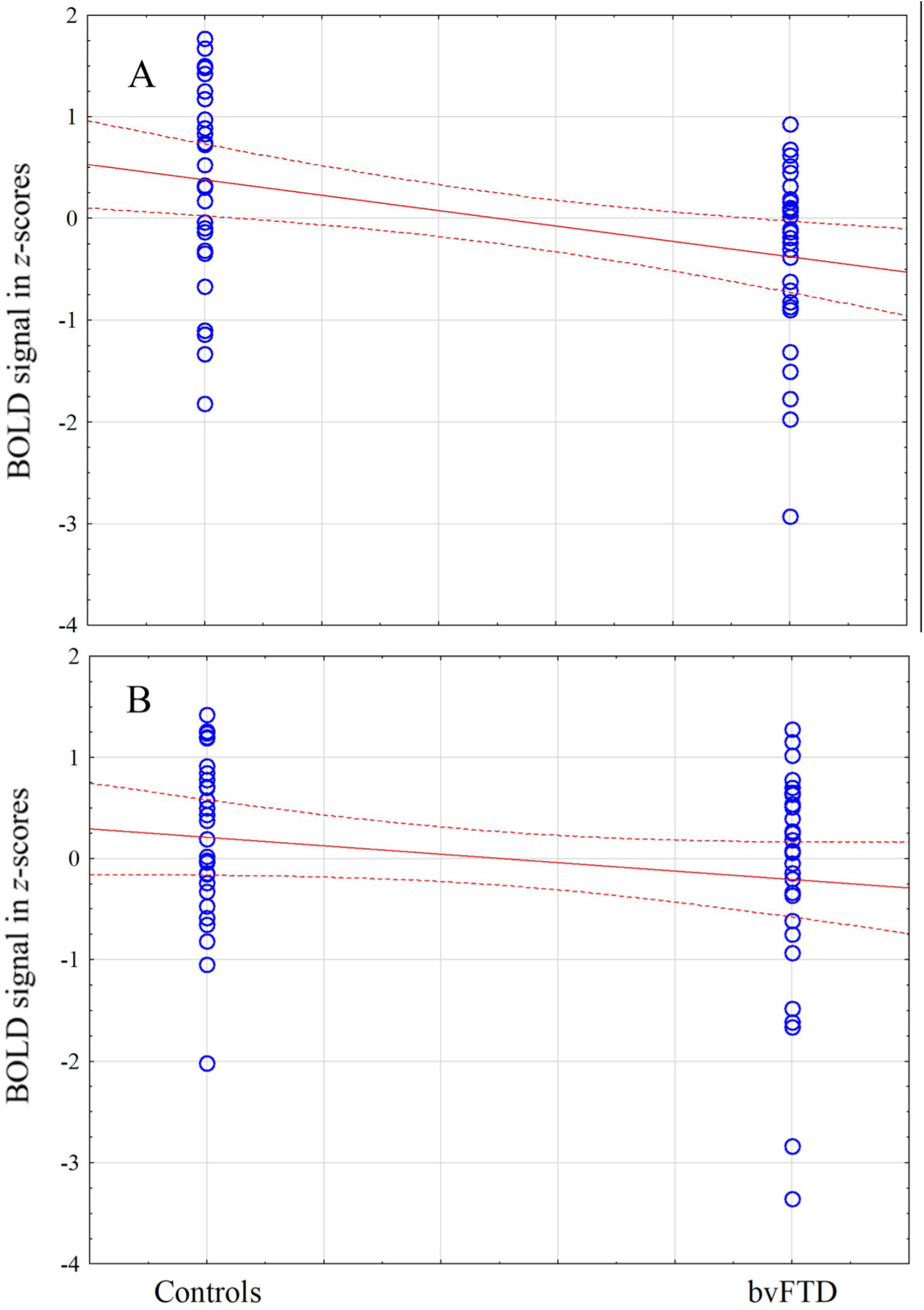
BOLD signal change in the empathy of pain contrast in patients and controls. This figure displays the percentage signal change, converted to *z-scores*, during the empathy of pain contrast under the affective perceptual empathy ROI (**A**) and the cognitive evaluative empathy ROI (**B**) in patients compared with controls. Blue circles indicate individual observations. Red solid line indicates mean. Red dotted lines indicate 95% confidence interval. Controls = control subjects; bvFTD = behavioural variant of frontotemporal dementia.

#### Relationship with the interpersonal index

BOLD signal under the CA-ROI in the EFP contrast was significantly positively correlated with the control subjects self-rating of their EC using the IRI (*r*=0.61, *p*<0.01), and with informants ratings of patients EC (*r=*0.50, *p*=0.03, Fig. 5). There were no significant correlations with the other IRI scales in the controls self-rating or the informants rating of patients (Supplementary Table 3). In contrast, patients own rating of their EC was not significantly correlated with their BOLD signal under the CA-ROI. There was no significant difference between controls and patients regarding self-rating on the IRI subscales. BOLD signal under the CA-ROI was not correlated with global cognition as measured with the MMSE (controls *r* = -0.26, *p* = 0.33, patients *r* = 0.06, *p* = 0.74) or with performance of any other neuropsychological test. Mean cortical thickness under the CA-ROI was not significantly correlated with self-rated (for controls) or informant rated (for patients) EC (controls, self *r* = 0.02, *p* = 0.94; patients, informants rating *r* = 0.40, *p* = 0.09). Finally, mean BOLD signal under two spheres located at peak BOLD signal in the EFP contrast in the patients (left occipital cortex and the left supramarginal gyrus) did not predict informants rated EC (*r =* 0.31, *p* = 0.19).

**Figure 5.**
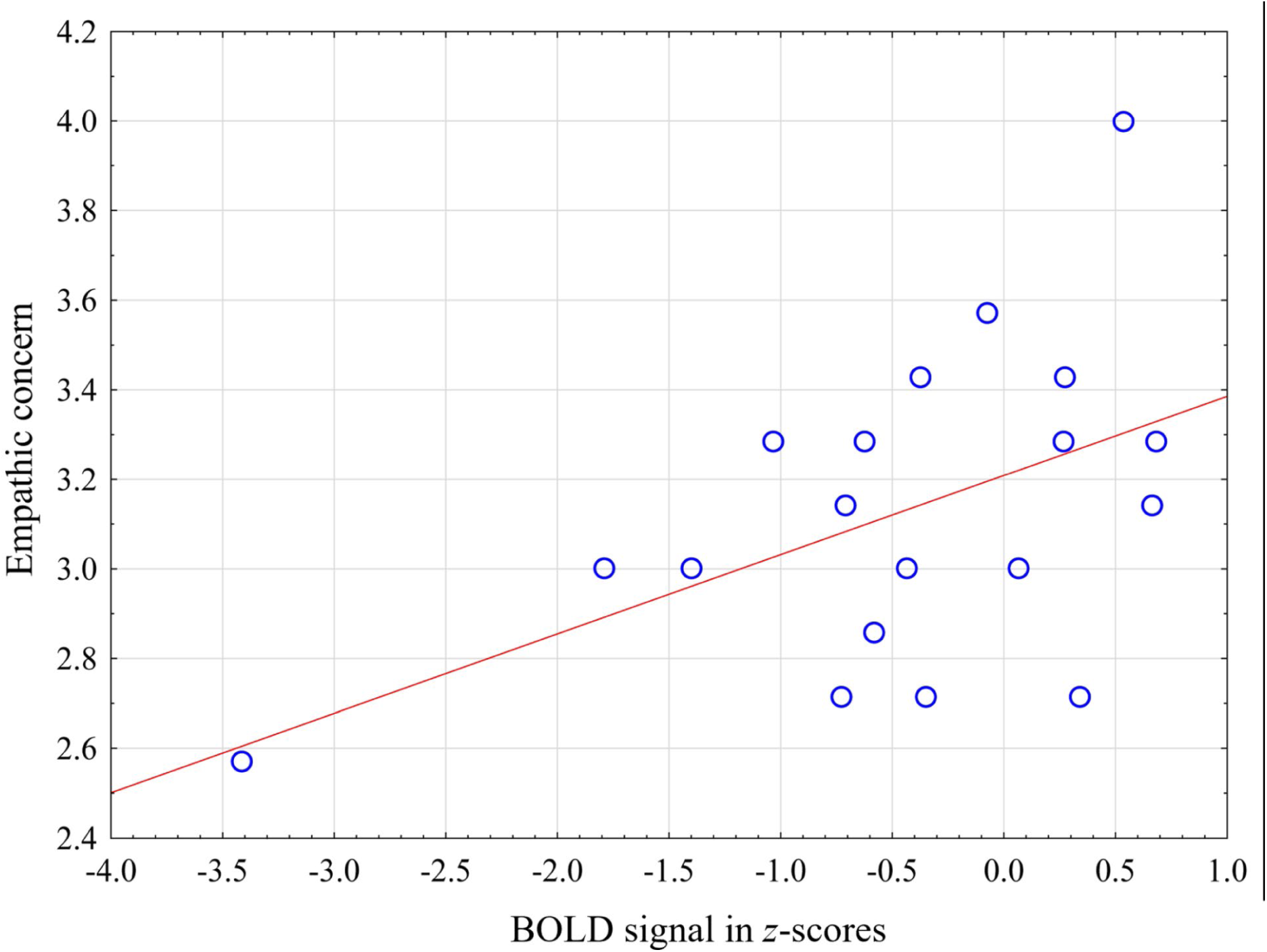
Correlation between empathic concern and BOLD signal in patients with bvFTD. The figure displays the correlation between mean BOLD signal in patients extracted under the control activation ROI (CA-ROI) and empathic concern in patients rated by informants. X-axis, denotes percent signal change during empathy for pain under the CA-ROI converted to *z*-scores. Y-axis denotes mean of empathic concern ratings in the IRI as rated by informants. BvFTD = the behavioural variant of frontotemporal dementia.

#### Seed-based resting-state functional connectivity analysis

In the ROI-to-ROI (sphere-to-sphere within the CA-ROI) correlation analysis we observed heightened synchronized fluctuations in the resting-state BOLD signal in controls compared to patients, especially in regions associated with empathy related activation (in the controls in the present study), also overlapping the VAN. For example, the resting-state BOLD signal in the right posterior orbitofrontal/insula had a significantly stronger correlation with the BOLD signal in the left insula, left dorsal anterior cingulate gyrus, and supramarginal gyrus in controls than in patients. However, the average BOLD signal correlations within the visual network, another key network in the EFP paradigm, showed no significant difference between the two groups (Fig. 6).

**Figure 6.**
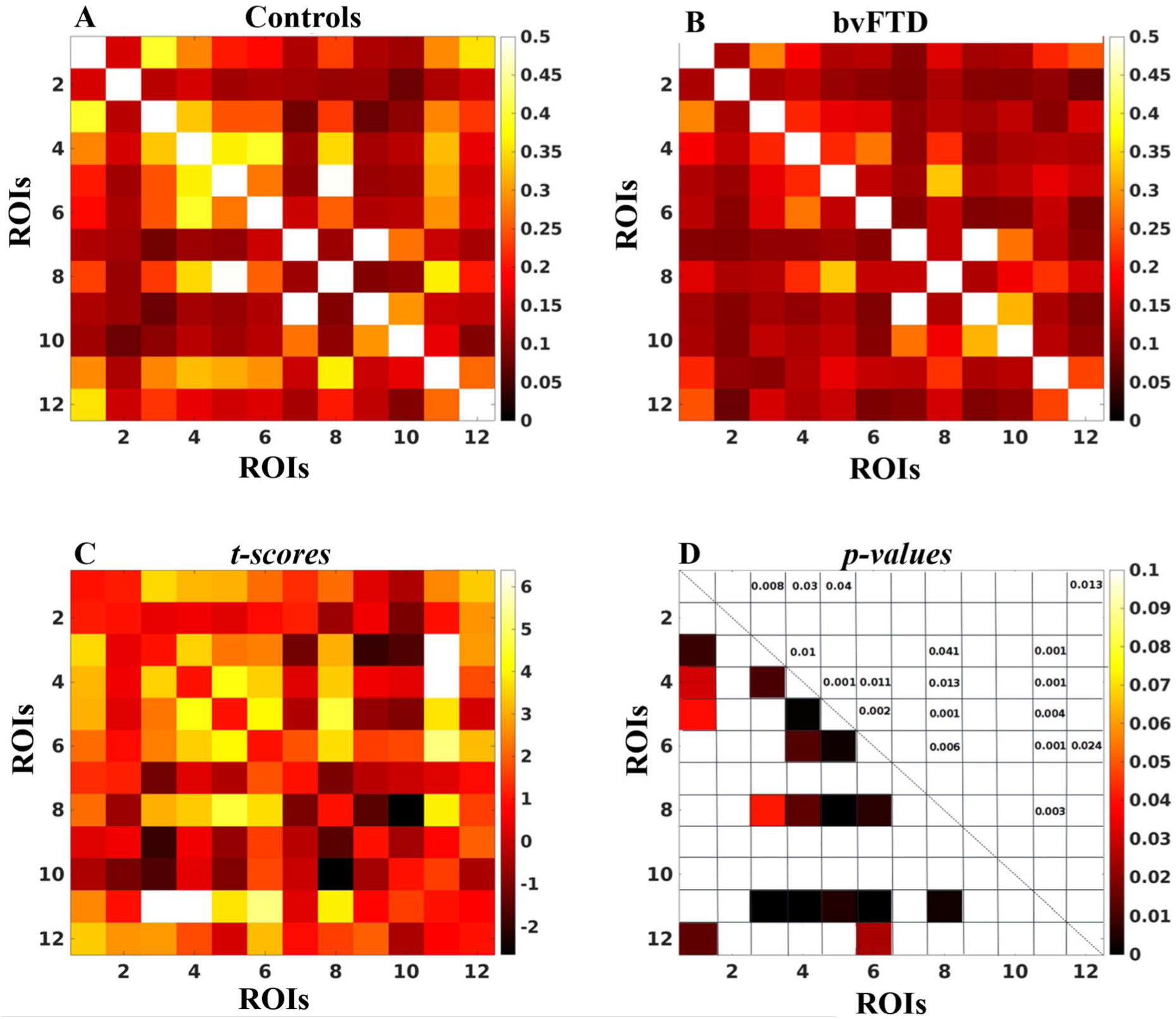
Resting-state connectivity between the 12 areas in the control activation ROI. Fig. 6A and **B** depict correlation matrices for average BOLD signals in each of the 12 seeds, corresponding to the 12 task defined ROIs in the control activation ROI, for the control and patient groups, respectively. *T-test* score differences between controls and bvFTD patients and corresponding *t-test p-values* are shown in Fig. 6C and **D** respectively. Area number in the matrices correspond to area numbers in Table 2.

In the 12 task-defined CA-ROI-to-all other voxels of the brain correlation analysis, we observed that areas overlapping the VAN displayed largest difference between patients and controls, less difference in the DMN and least in the visual network. Stronger correlation in controls were mainly observed in areas belonging to the same network as the task-defined ROI (Supplementary material Fig. 6.1-6.12, & Resting_state_Supplementary_table.xlsx).

#### Diagnostic performance of BOLD signal

BOLD signal in the EFP contrast under the was a modest discriminator between patients and controls. A receiver operating characteristics curve (ROC) analysis revealed an area under the curve (*AUC*) of 0.74, (Supplementary Fig. 7), using the mean BOLD signal under the affective perceptual empathy ROI as a test variable. The mean thickness of the right insula was a better discriminator, with an *AUC* of 0.86, (Supplementary Fig. 8).

## Discussion

We show in a well-established task-based fMRI paradigm for EFP that bvFTD patients display a different activation pattern than healthy individuals, predominantly in regions that are characteristic targets of bvFTD related neuropathology. Differences were observed in regions known to be important for the ability to experience affective perceptual but not cognitive evaluative empathy. Strength of activation was related to a measure of affective empathy in real life. The intrinsic resting-state connectivity between regions associated with empathy processing during task-based fMRI was further observed to be weaker in patients than in controls, predominantly in regions overlapping the VAN. To the best of our knowledge this study provides a missing link between on one hand a core bvFTD symptom and on the other hand the characteristic atrophy pattern of this condition and the deficit in the corresponding intrinsically connected network.

As hypothesized, empathy for pain was associated with increased BOLD signal in the AI and the ACC/MCC in controls, areas that have been shown to be among the earliest affected in bvFTD.^14,60^ Patients showed no significantly increased activation in these regions during EFP. Furthermore, the patients displayed significantly reduced BOLD signal in regions related to affective perceptual, but not cognitive evaluative, aspects of empathy processing, in line with the literature showing predominantly affective empathy deficits in bvFTD.^5–8^ We observed a common activation in both patients and controls occipital cortex and left supramarginal gyrus. The latter finding is in line with previous meta-analysis that showed activations in this area in picture-based EFP paradigms.^39^ One interpretation is that it is the aspect of intentionality attribution of the cue-based paradigms (and the role of the supramarginal gyrus in these functions) that leads to this pattern.^61^ The supramarginal gyrus is furthermore part of the VAN,^19^ and was atrophic in the patient group, as depicted in Fig. 2A. Interestingly, cortical thickness in left supramarginal gyrus in patients was not correlated with the strength of BOLD signal in this area (Pearson’s *r =* 0.0 *p* = 0.97).

As a measure of the participants ability to experience empathy we used self-ratings (in controls and patients) and informant ratings (in patients) of the IRI subscales. The CA-ROI was used as it reflected the expected normal activation pattern of the specific empathy task used in this study. While the analysis in controls was merely used to confirm validity, the main analyses was performed in the patients. Notably only the EC subscale displayed significant correlation with BOLD signal in the EFP contrast, in both patients (when rated by an informant) and controls (when rated by themselves). In contrast to the ratings by informants, the patient ratings of empathy did not relate to the empathy associated activation. Our interpretation is that this is due to lack of insight that is a common symptom in bvFTD.

Studies on healthy volunteers have demonstrated a task-dependent laterality effect in empathy associated activation of the AI. Singer et al. used abstract cues to indicate when a painful electric stimulation was administrated to another person’s hand. They found that percent signal change in the left AI during EFP cues was positively correlated EC in IRI.^37^ Li et al. found that percent signal change in the right AI during affective perceptual empathy was positively correlated with EC in IRI.^62^ In contrast to findings by Li et al. and Singer et al., we did not find a significant correlation between BOLD signal in neither left of right AI and any IRI subscale, but instead with the mean BOLD signal in all regions activated during EFP in controls. Li et al. proposes that people with high affective empathic ability have sensory system that are more active than people with low affective ability,^62^ which is in line with our finding that we see more activity in the whole CA-ROI in participants with high EC-scores.

While we did not find significant correlation between right nor left AI BOLD signal and informants rating of EC in patients, we still found that right but not left AI (red spheres, Supplementary Fig. 2) displayed significantly decreased BOLD signal in patients compared to controls in the during EFP (Supplementary Fig. 9A [right] Fig. 9B [left]). We propose that the difference in the right but not left AI may be explained by two different factors. First, it is known that affective perceptual empathy involves the right AI more than the left.^40,62^ Second it is known that bvFTD-pathologies selectively causes neurodegeneration in a specific cell type: The von Economo neurons (VEN) that are located in the ventral AI and the ACC.^63^ Interestingly there are significantly more VEN in the right than the left hemisphere,^63^ which may explain why several studies observed more atrophy in the right than left insula in bvFTD,^14^ (see Schroeter et al,^64^ for meta-analysis). Thus, independent of fMRI task, it is likely that we may observe more functional and structural changes in the right AI than the left AI in patients with bvFTD.

It should be noted that EC, synonymous with compassion,^32^ is not only involving the ability to experience the emotional state of another person but also as a strong motivation to improve the other’s wellbeing.^31^ The later aspect of EC has been associated with activation of the braińs reward circuit involving regions such as the medial orbitofrontal cortex, nucleus accumbens & subgenual ACC.^31,32^ However, importantly, also regions such as insula and caudal ACC (that showed attenuated activation in patients) are activated in EC and provide a support function for the second order reward processes in compassion associated activity. Studies in bvFTD have attempted to disentangle these two aspects of EC, and indeed demonstrated that it is the affect sharing part of EC that is related to insula ACC, while prosocial behaviour is linked (as in healthy controls) to a reward networks.^65^

An noteworthy finding was that functional BOLD signal increase during EFP but not structural changes (mean cortical thickness) under the CA-ROI, was associated with patient’s ability to experience EC. Further, neither BOLD signal during EFP nor EC ratings in patients, was correlated with patients’ performance on the other neuropsychological tests available in this study. Thus, that BOLD signal during EFP was the only imaging or neuropsychological data that was associated with EC is in line with a previous finding showing that EC in the IRI reflects unique characteristics of emotional information processing that seem unrelated to abilities that are assessed with a traditional neuropsychological battery.^5^

We found largest difference between patients and controls in synchronized resting-state fluctuation of BOLD signal in areas overlapping the VAN in the CA-ROI. Less difference was observed for the DMN and least for areas belonging to the visual network. Some authors have suggested that the VAN may inhibit neural activity in the DMN. Deficits in the VAN could thus lead to an upregulation of the DMN.^20^ However, we did not observe a stronger correlation between the seed in the posterior cingulate cortex and other parts of the DMN in patients than in controls, which is in line with a recent review that found inconsistent evidence for this hypothesise.^18^

Previous studies offer several models for how the functional deficits in the VAN may affect patients’ capacity to experience empathy. Ibanez et al. propose a model for a “social context network”, which processes “social context effects”,^66^ suggesting that most cortical regions that become atrophic in bvFTD are involved in the social context network: the prefrontal cortex makes prediction based on context, the anterior insular cortex coordinates interoceptive and exteroceptive processing, and the anterior temporal cortex is involved in context-dependent associative learning. Deficits in social cognition in bvFTD, can with the social context network- framework be described as a deficit in the ability to process relevant cues in a social context (for example facial expressions in people during social interaction). Thus, patients may not fail to react emotionally because they lack the capacity to react, instead they fail to interpret emotional content of the situation.

Other models centre around the hypothesis that bvFTD patients have a diminished capacity for perceiving moment to moment changes in the physiological condition of the body, synonymous with interception.^67^ For example, Carlino et al. showed that patients with bvFTD had a higher threshold and higher tolerance for the experience of pain compared to healthy volunteers,^68^ which is consistent with a model that holds that the ACC,^69^ and the insula,^70^ have important roles in pain perception^43^. Interception deficits have indeed been demonstrated in bvFTD and shown to be related to social cognitive function.^71,72^

Damasio and Carvalho propose that feelings are mental experiences of body states, which are “directly portraying the advantageous or disadvantageous nature of a physiological situation.^73^” Feelings may, however, also occur as an efferent event in which visceromotor responses provide “a efference copy to alert the afferent division of the salience network.^20^”

In our experiment it is possible that the sight of a needle in a hand, simulating a needle in the participant’s own hand, may induce a rapid immediate response in the ACC to allocate physiological bodily resources to withdraw the hand. Information processing in the afferent division of the salience network may then make these bodily changes perceivable as feelings. The common activation of the left supramarginal gyrus in both patients and controls support such an interpretation, as previous studies have shown that this region, particularly on the left side is involved in action planning (e.g. moving the hand away from the needle,^74,75^). Notably, the social context network model may not be contradictory to models focusing on interoception. In line with Damasio’s somatic marker model, it could be hypothesized that patients with bvFTD lack the support of “somatic marker” (or interoceptive signal) in complex social situations, which may be the ultimate cause for why the social context network is failing to focus on and perceive the most relevant social cues of that situation. The discussion above potentially implies two ways of interpreting the diminished response during EFP in the patients. One possibility is that patients fail to focus attention to the salient cue (the needle in the hand) in the pain condition, due to lack of support from interoceptive signals/somatic markers. Another possibility is that they fail to experience discomfort at the sight of the needle in the hand because neurodegeneration has occurred at the centre (the AI) in which interoceptive and sensory signals are integrated and perceived as a “global emotional moment.^22^” These two explanations are not mutually exclusive. In fact, there may be an interaction between them, in which the lack of attention to salient stimuli reduces the emotional reaction, and the reduced emotional reaction further reduces attention to salient stimuli.

Another aim of this study was to investigate whether BOLD signal in the empathy for pain paradigm would be useful for diagnostic purposes. BOLD signal was, however, only a modest discriminator between patients and controls, with less accuracy than structural changes e.g., the mean thickness of the right insula. Thus, the paradigm is more useful for predicting the capacity to experience empathy than for diagnosis. While we emphasized the fact that loss of empathy is a central symptom in bvFTD, it is also known that the predominant symptoms at early stages may differ between individuals,^2^ and it is possible that such variation in our sample may explain the poor results in the ROC analysis with BOLD signal as test variable.

A limitation of this study is that three different MRI scanners were used, and while there was no significant difference in BOLD signal in the EFP contrast between patients investigated in the different scanners, some scanner-related variation in the BOLD signal that we are not able to detect is expected. We therefore performed a separate analysis including only patients (n=21) and controls (n=13) investigated in the Prisma MRI scanner in Lund. This analysis replicated the results from the whole cohort both in the reduced BOLD signal in patients compared to control under the affective perceptual empathy ROI, and the significant association between mean BOLD signal and informants rating of patient’s empathic concern ( Supplementary sensitivity analysis 3&4 and Supplementary Fig. 4). Thus, we conclude that MRI scanner is not a problematic confounder in this study. Considering the significant association between BOLD signal in our fMRI paradigm and EC from the IRI, it would have been interesting to see whether BOLD signal also would be linked to other tests assessing changes in the patient’s personality or measurements of other aspects of the EC such as compassion/motivation.

In conclusion, patients with bvFTD displayed decreased empathy related neural activity in regions including the bilateral anterior insula and the ACC, which have previously been shown to be of central importance for empathy processing and are known to be affected by pathology early in bvFTD. The magnitude of empathy related neural activity predicted patient’s ability to experience affective empathy. This shows that indeed patients with bvFTD have altered empathy processing, bringing together, and consolidating previous empirical evidence, demonstrating a state-dependent functional change in core bvFTD networks, that is associated with a core clinical symptom in this disease.

## Data availability

Anonymized data will be shared by request from a qualified academic investigator for the sole purpose of replicating procedures and results presented in the article if data transfer is in agreement with relevant legislation on the general data protection regulation and decisions and by the relevant Ethical Review Boards, which should be regulated in a material transfer agreement.

## Supplementary material

Supplementary material is available at *Brain* online.

## Authors contributions

OL, GN, PP, LW, AS, LN: conceptualization, study design, formal analysis, writing - original draft.

OL: was responsible for coordinating data collection at the three different sites, scanned patients in Stockholm, and performed the analyses of structural and functional imaging data.

LW: the project was originally initiated by LW who supervised several previous studies on empathy in various cohorts for example patients with psychopathy.

GN: provided the original proposal for how this paradigm could be used in the study, and participated in detailed planning of the design of the study.

GN, PP: have previously used the EFP-paradigm in several fMRI studies and provided OL with crucial support during the analysis and the interpretation of the imaging data in the study.

LN: have provided important input on how fMRI data should be collected and analysed. Further he provided crucial support on how results of the study should be interpreted.

AS: harmonizing the diagnostic procedures and diagnostic quality control.

TL: performed analysis of resting-state data.

CL, NB, LW, AS, OH, CC: recruited and diagnosed patient.

SV, OA, AE: performed neuropsychological testing of patients.

OA was further responsible for creating a final database for test-scores for patients recruited at different centres.

PM: was MRI physicist in the project.

MS: edited the images used in the fMRI paradigm and collected IRI-rating for patients in Umeå.

LH: manuscript writing, editing and review.

All authors reviewed and edited the manuscript and approved the final version.

## Supplementary material

### The Swedish text that are displayed during the fMRI paradigm

The text “What is the hand feeling” is in the Swedish version used in this paradigm translated into “Vad känner handen?”.

### Acquisition and analyses of structural MRI

#### Structural MRI

In the Prisma MRI scanners structural 3-D T1-weighted images were acquired with a voxel size= 1x1x1.2 mm^3^, with an inversion time (TI)=900 milli seconds (ms), repetition time (TR)=7100 ms, echo time (TE)=2.98 ms and a flip angle (FA) of 9 degrees. In the GE-MRI scanner the T1was acquired with a SPGR 3-D sequence with a TR=8156 ms TE =3.18, TI=450 ms, a FA 12 degrees, and a voxel dimension of 1x1x1mm^3^.

#### Resting-state MRI

In the Prisma MRI scanners resting-state fMRI images were acquired with a TR/TE=2030/30 ms, , with an in-plane resolution of 2.5*2.5 and slice thickness of 3.8 mm and FA=80 degrees. In the GE MRI scanner resting-state fMRI images were acquired with a TR/TE=2000/30 ms, , with an in-plane resolution of 1.9*1.9 mm and slice thickness of 3.8 mm. The total acquisition time for the resting-state fMRI was consistently 8 min.

### Image quality control

Quality control was carried out on all MRI data according to previous described procedures ^1^, and data management and processing were done through our in house database system ^2^. As the registration process is particularly challenging in patients with significant brain atrophy, this process was visually inspected for each subject included in the study (which can be reviewed in supplementary material: Registration_All.pdf). During the experimental fMRI acquisition, all patients and controls had a relative displacement of less than a half voxel (>1.15 mm) and an absolute displacement of less than a voxel (>2.33 mm). The mean relative movements (between one acquired volume to another) in controls was 0.08 mm and in patients 0.11 mm. This difference was not significant *(p* = 0.07),

#### Structural MRI analysis in FreeSurfer

Cortical reconstruction and volumetric segmentation of subcortical volumes were performed on T1 3D images using <u>Freesurfer</u> 6.0.0 image analysis pipeline, which is documented and freely available for download online (http://surfer.nmr.mgh.harvard.edu/). The technical details of these procedures are described in prior publications, which are listed at https://surfer.nmr.mgh.harvard.edu/fswiki/FreeSurferMethodsCitation. Briefly, the whole-brain T1-weighted images underwent a correction for intensity homogeneity, skull striping, and segmentation into GM and white matter (WM). Cortical thickness was measured as the distance from the gray/white matter boundary to the corresponding pial surface. Subcortical segmentation and assessment of intracranial volume was also performed in Freesurfer. Reconstructed data sets were visually inspected for accuracy, and segmentation errors were corrected.

#### Statisticalanalysis

Group comparisons in cortical thickness measures was performed using vertex-based GLM (general linear model) in the FreeSurfer software correcting for the effect of gender. The Gaussian smoothing kernel was 10 mm. The level of statistical significance was evaluated using a cluster-wise *P* (*CWP*) value correction procedure for multiple comparisons based on a Monte Carlo z-field simulation with a cluster forming threshold of *p* of cluster > 0.01 (vertex- *z*-threshold = 2).

### Analysis of resting-state functional MRI

#### Preprocessing

The R-fMRI datasets were preprocessed using AFNI (Version Debian-16.2.07dfsg.1- 3nd14.04+1, link) and FSL (link) tools ^4^. Procedures involved temporal de-spiking, six-parameter rigid body image registration for head-motion correction, and generation of a brain mask from the average volume of motion-corrected series to exclude extra-cerebral tissues. Spatial normalization to the MNI standard template utilized a 12-parameter affine transformation with a mutual-information cost function, and data was re-sampled to isotropic resolution with a 4 mm FWHM Gaussian kernel ^5,6^. The cohort’s average image volume was used to create an average brain mask. Nuisance signals were removed using multiple regressors for motion correction parameters, ventricle signals, and their derivatives. After removing baseline trends up to third-order polynomial, band-pass filtering was applied at 0.08 Hz. Local Gaussian smoothing was conducted up to FWHM = 4mm using an eroded gray matter mask ^5^.

#### Statistical Analyses

Connectivity patterns of brain regions that are activated during the EOP task were assessed using Pearson’s correlation coefficients (CC) between the average time courses of the 12 task-defined ROIs, creating a 12x12 symmetric matrix per subject. Differences between patients and controls were evaluated using a two-sample t-test on these matrices. The Bonferroni approach was employed for multiple comparison correction.

To explore the intrinsic connectivity network of each of the 12 task-defined ROIs and any bvFTD pathology impacts, Pearson CC maps were computed between each seed ROI’s average time course and every resting-state-fMRI data voxel for all 12 seed ROIs. To distinguish connectivity differences between bvFTD patients and controls, two sample t-tests were performed on these maps. A two-step approach was used for statistical significance. First, a voxel-wise threshold of *p*<0.001 (*t*-score ≥3.3) was set for initial cluster candidates. Then, permutation simulations identified significant brain regions from initial clusters at a family-wise error rate (FWER) of *p*≤0.05.

### MRI-scanners

Quantification of effect size of the response under the affective perceptual empathy ROI in patients investigated with the three different MRI scanner revealed no significant difference: Coheń s *d* (MRI scanner 1 vs. MRI scanner 2) = 0.86; *p* = 0.41, MRI scanner 1 vs. MRI scanner 3, *d* = 0.32; *p* = 0.51; (MRI scanner 2 vs. MRI scanner 3), *d* = 0.38; *p* = 0.74.

### Sensitivity analyses for the task-based fMRI

In sensitivity analysis one, we investigated difference between patients and controls under the affective perceptual empathy ROI in the whole sample correcting for mean relative displacement. Difference between controls and patient remained significant [Current effect: *F*(1,53)=6.5, *P*<0.05].

In sensitivity analysis two we included participants that were investigated in the Siemens Prisma cameras. In the model we included diagnosis, sex and site (KI camera vs. Lund camera) as categorical predictors, age as continuous predictor and BOLD signal under the affective perceptual empathy ROI as dependent variable. Difference between controls and patient remained significant [Current effect: *F*(1,46)=5.6, *P*<0.05].

In sensitivity analysis three we included participants that were investigated in the Prisma Camera in Lund. Diagnosis and sex were included as categorical variables, age as continuous predictor, and BOLD signal under the affective perceptual empathy ROI as dependent variable. Difference between controls and patient remained significant [Current effect: *F*(1, 30)=4.6*, p*<0.05].

In sensitivity analysis four we investigated the association between BOLD signal under the affective perceptual empathy ROI during EFP and informants rating of patients EC including only participants investigated in the Siemens Prisma camera in Lund. The correlation remained significant (*r* = 0.50, *p*<0.05). Supplementary Figure 4 below.

In sensitivity analysis five we investigated the association between cortical thickness under the control activation ROI and BOLD signal during EFP in patients and controls. This correlation was not significant and very week (approximately *r*=0.1; Supplementary Figure 5 below). The results remained basically the same if BOLD-signal was correlated with mean thickness of the whole left and right hemispheres.

---

**Figure S1.**
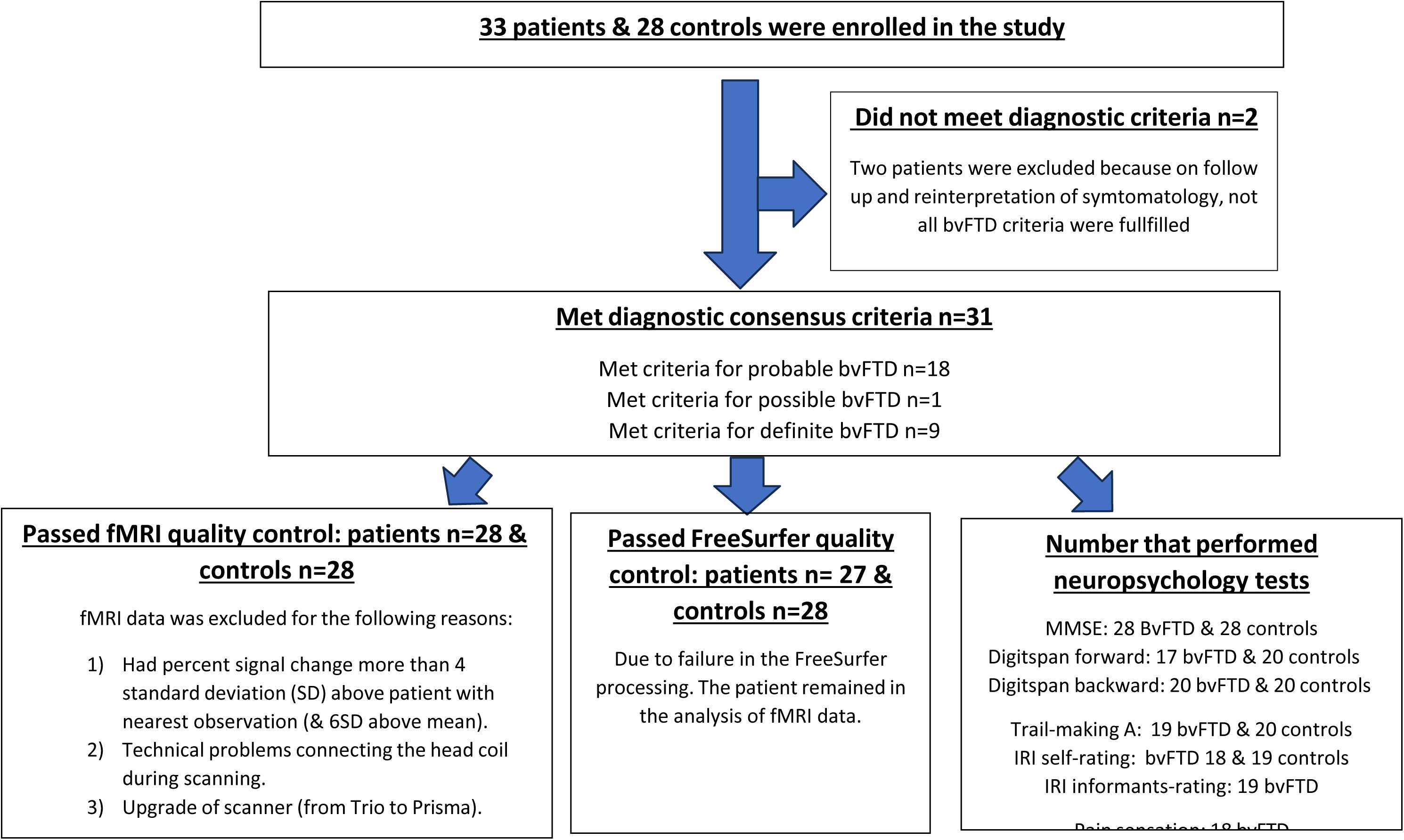
Flowchart displaying inclusion procedures of patients and controls.

**Figure S2.**
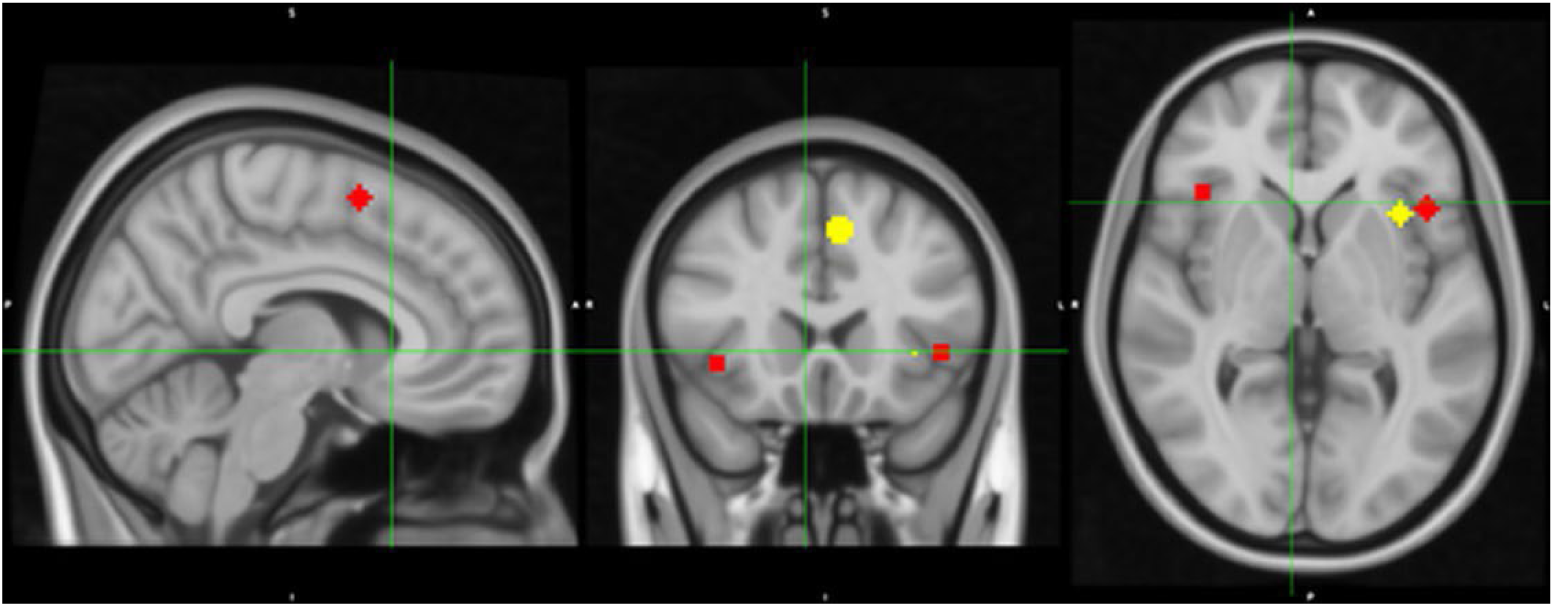
Spheres located at coordinates with peak activation in affective perceptual & cognitive evaluative empathy in the meta-analysis by Fan et al. Red spheres (the Affective perceptual empathy ROI) located at peak activation in affective empathy. Yellow spheres (the cognitive evaluative empathy ROI) located at peak activation for cognitive evaluative empathy in the meta-analysis, table 2 in Fan et al. 2011 ^3^

**Figure S3.**
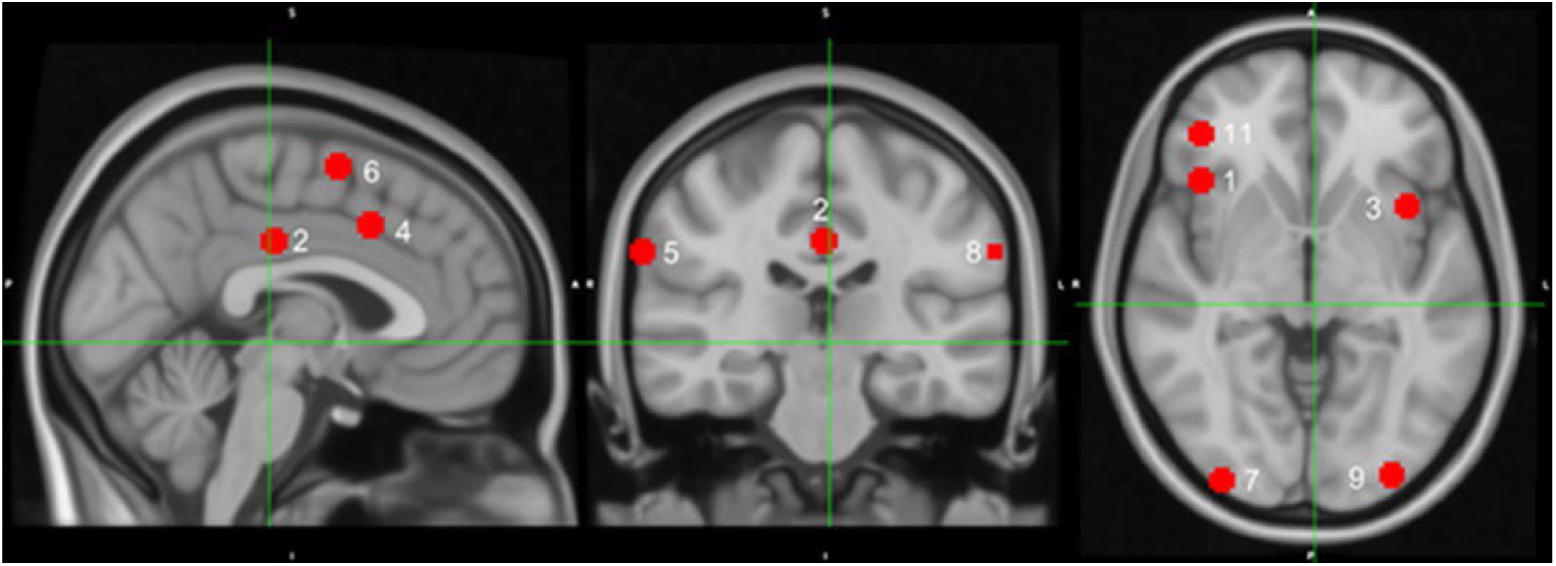
12 task-defined ROIs based on the significant activation in the empathy of pain contrast in controls. The figure displays 12 task defined 5mm spherical ROIs located at peak activation in the 12 areas with significant activation in empathy of pain in controls. Mean BOLD-signal was extracted for the 12 ROIs (the CA-ROI). The sphere located in the right frontal pole (number 12), is not visible at image above.

**Figure S4.**
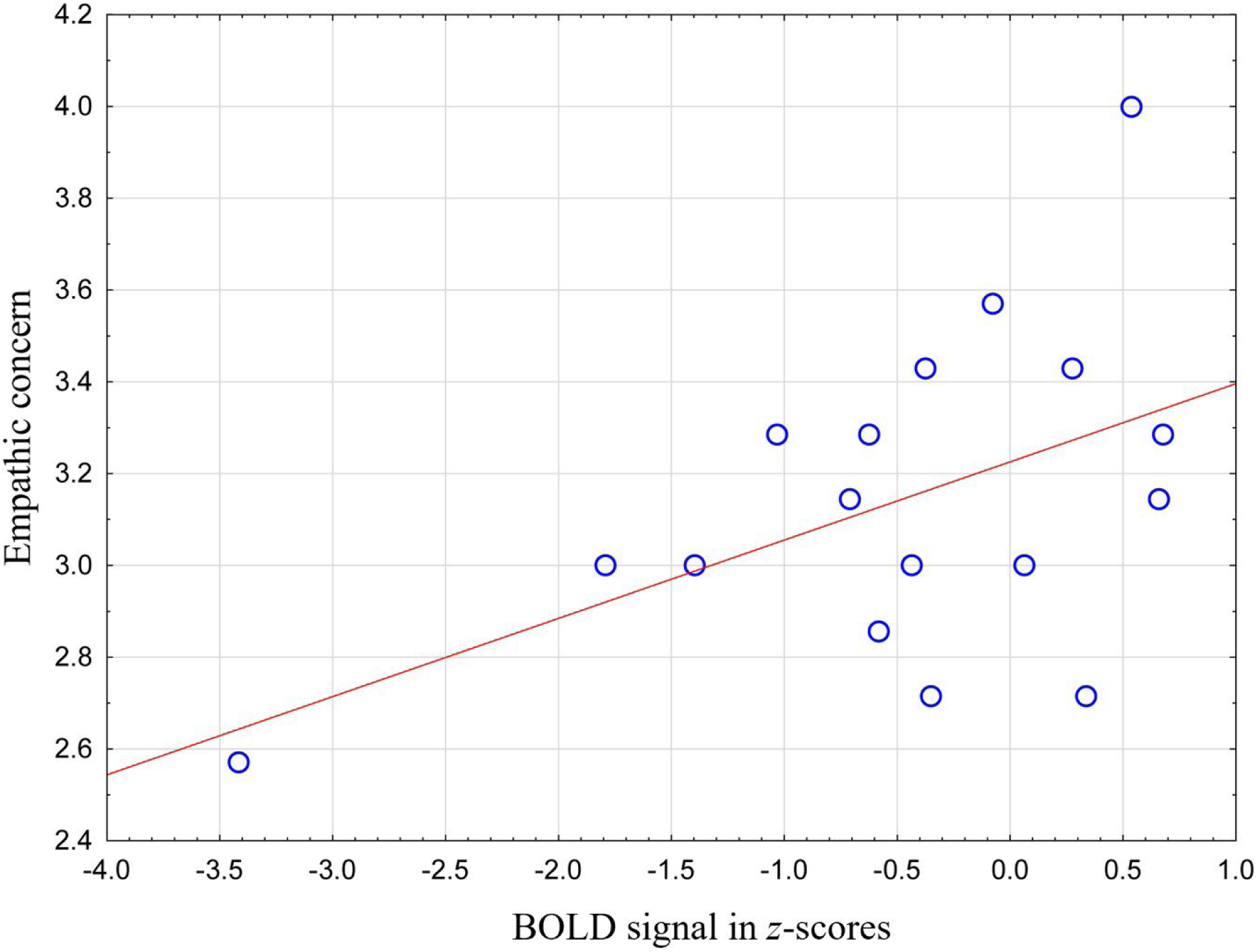
The correlation between informants rating of EC and BOLD- signal under the affective perceptual empathy ROI in patients investigated in the Lund camera. The association between EC and BOLD-signal under the control activation ROI including only participant investigated in the Lund camera. Blue circles represent individual subjects. X-axis, denotes percent signal change during empathy for pain under the CA-ROI converted to *z*-scores. Y-axis denotes mean of empathic concern ratings in the IRI as rated by informants. BvFTD = the behavioural variant of frontotemporal dementia.

**Figure S5.**
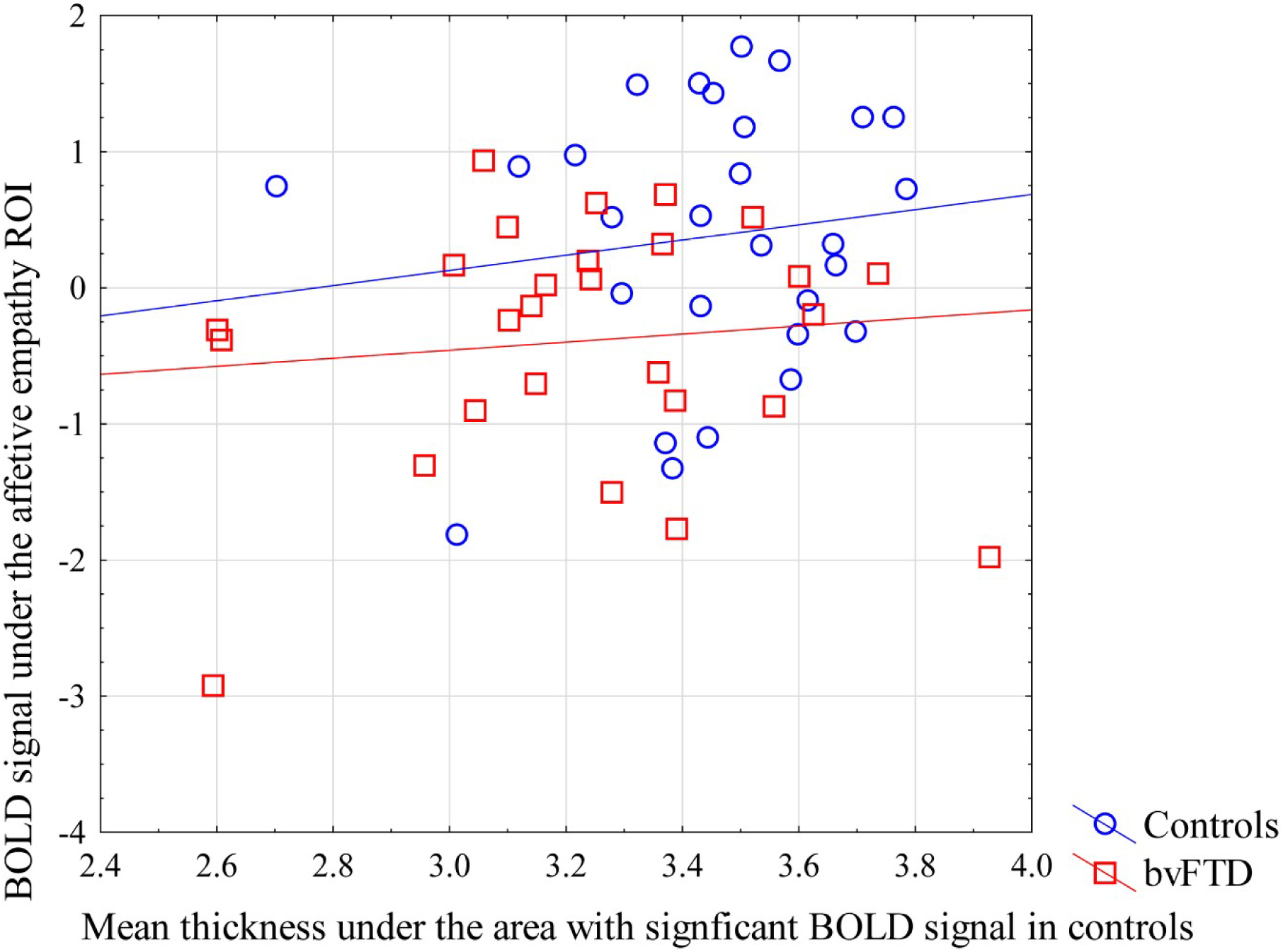
The correlation between cortical thickness and BOLD-signal under the affective perceptual empathy of pain ROI. The Graph is displaying the correlation between BOLD-signal under the affective perceptual empathy ROI and cortical thickness under the area with significant BOLD-signal in controls.

**Figure S6.**
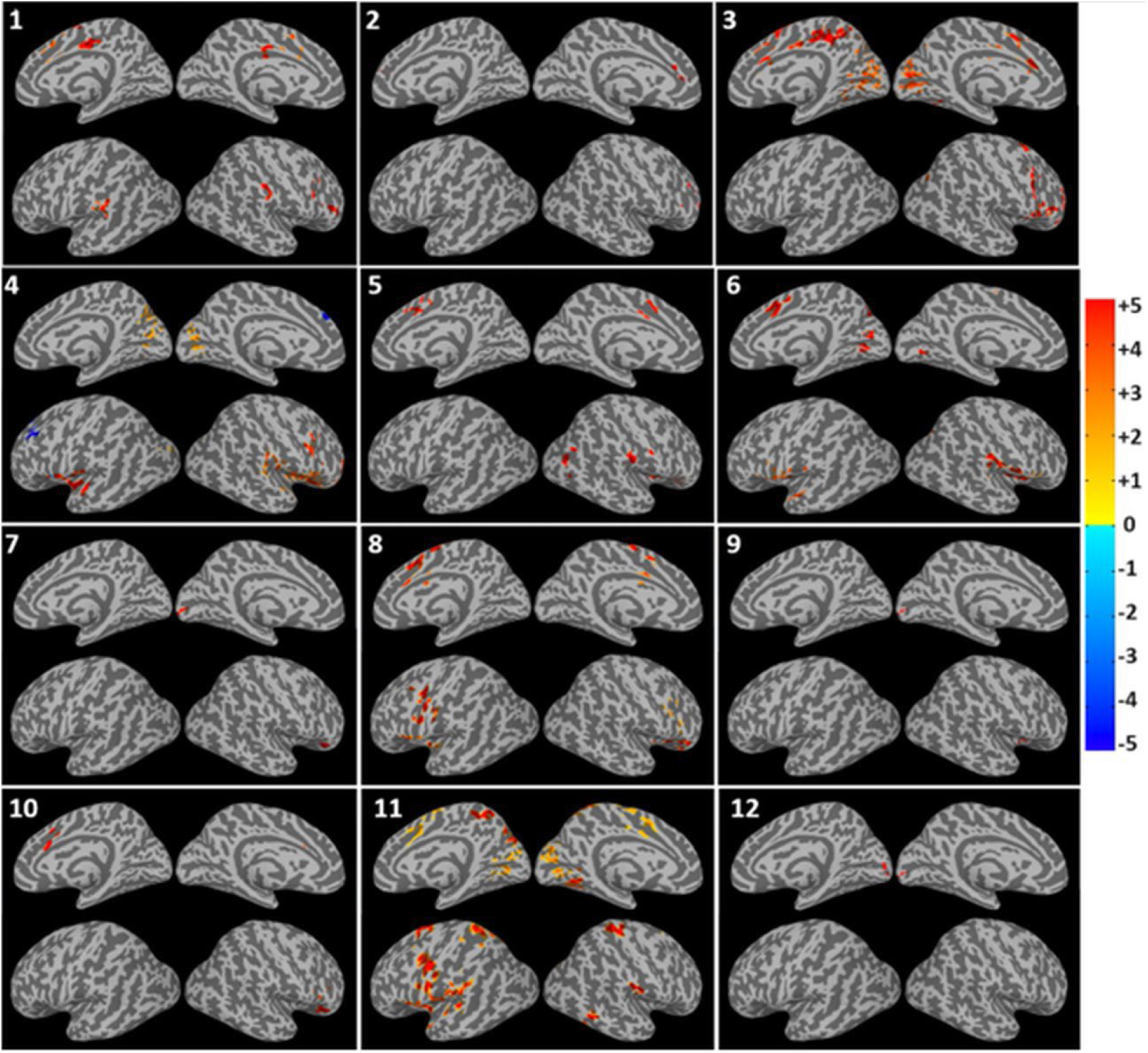
Stronger seed-to-whole-brain correlation of resting-state BOLD- signal in controls compared to patients. The figure depicts the difference between patients and controls in the correlation between the mean time-series for each of the 12 seed (task-defined ROIs) in the control activation ROI and every other voxel of the brain. Seeds: 10.1, right frontoinsular area, 10.2, bilat posterior cingulate, 10.3, left insular cortex, 10.4, left anterior cingulate, 10.5, left supramarginal gyrus, 10.6, left supplementary motor area, 10.7, right occipital pole, 10.8, right supramarginal gyrus, 10.9, left occipital pole, 10.10, left occipital fusiform gyrus, 10.11, right inferior frontal gyrus pars opercularis, 10.12, right frontal pole. Warmer colors indicate stronger correlation in controls. Cold colors (only ROI 4) indicate stronger correlation in patients.

**Figure S7.**
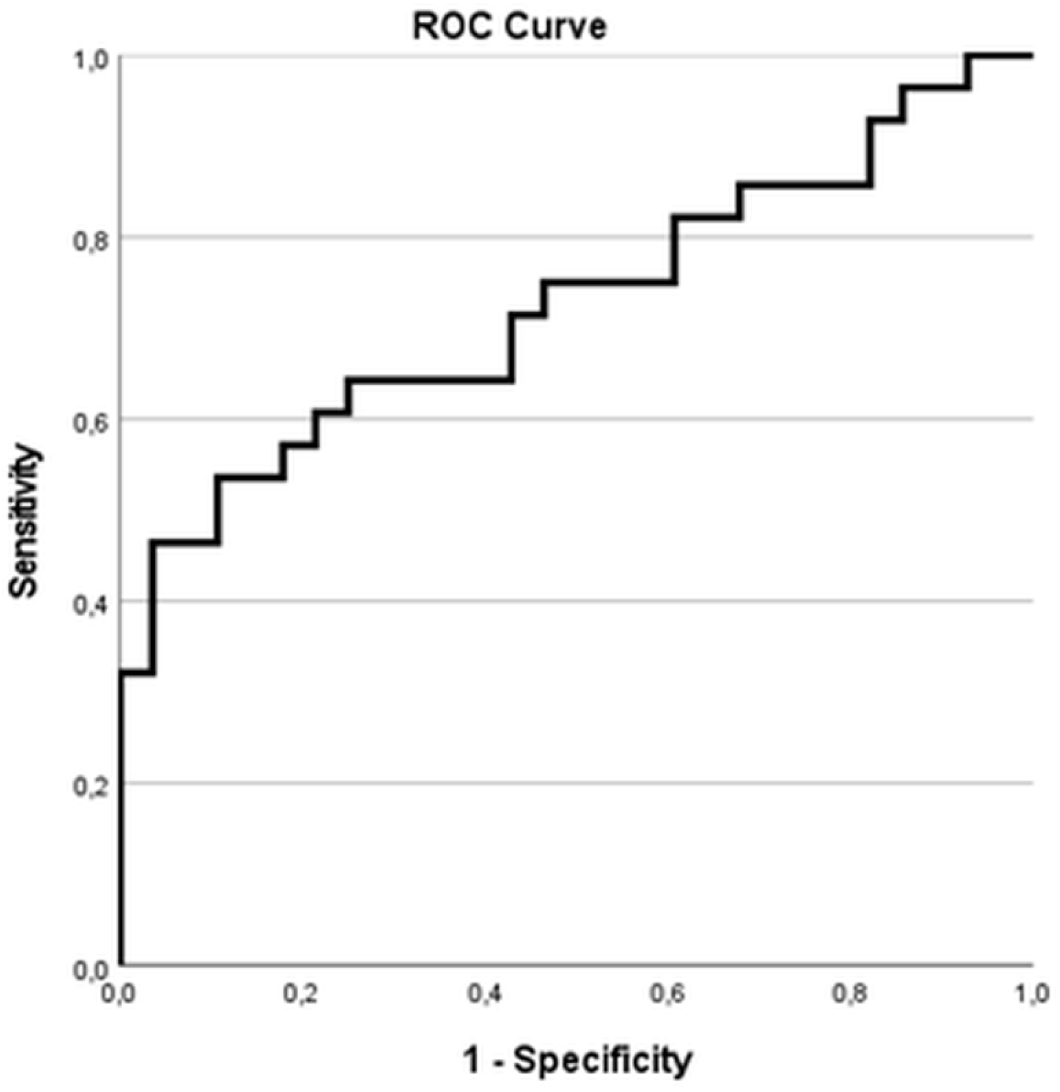
ROC-curve analysis with BOLD-signal under the affective perceptual empathyROI, during EP, as test variable. Results reveal an area under the curve = 0.73, SD = 0.069, Asymptotic Sig = 0.001; 95% Confidence interval 0.592 – 0.862.

**Figure S8.**
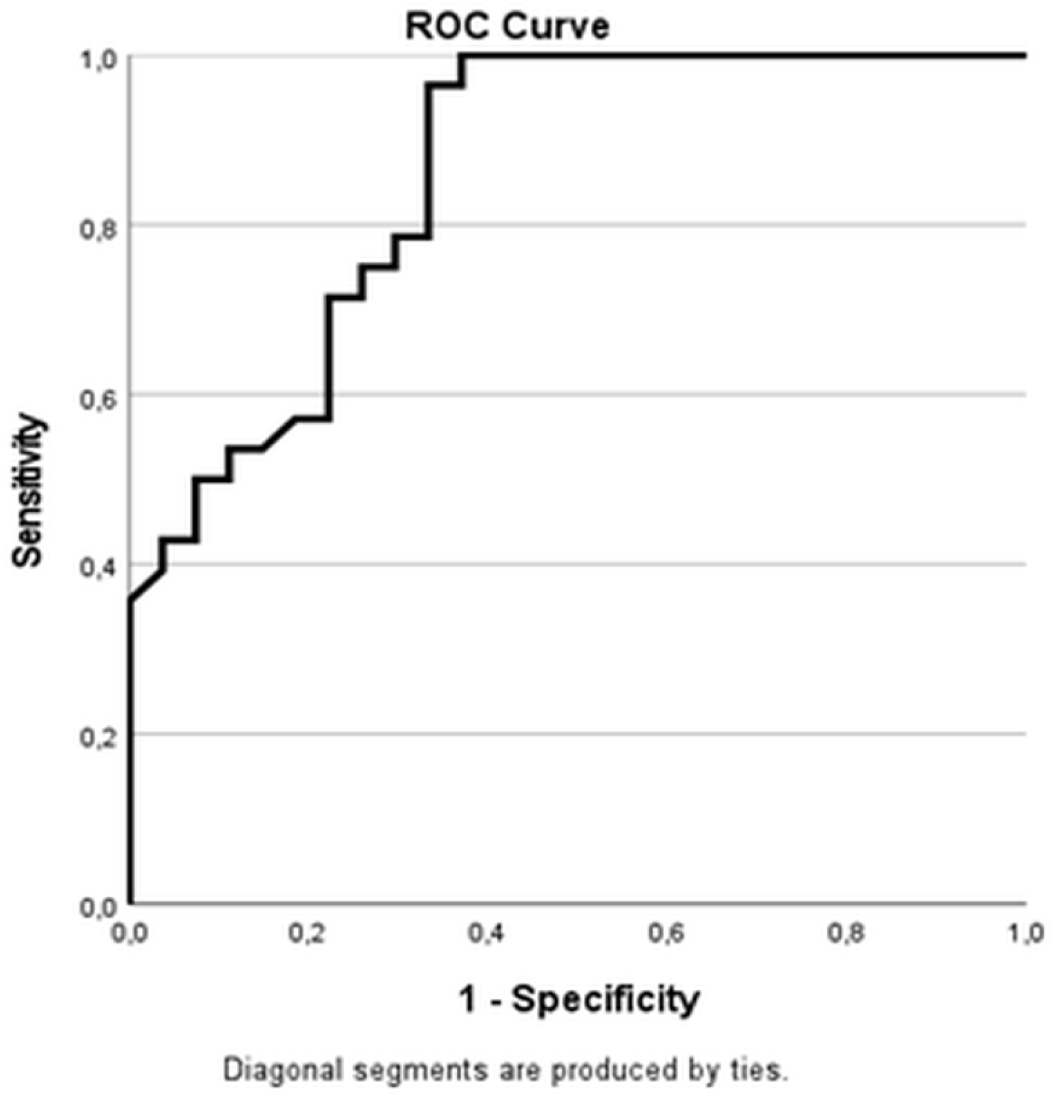
ROC-curve analysis with cortical thickness in the right insula as test variable. Results reveal an area under the curve = 0.858, SD = 0.049, Asymptotic Sig = 0.000; 95% Confidence interval 0.762 – 0.955.

**Figure S9.**
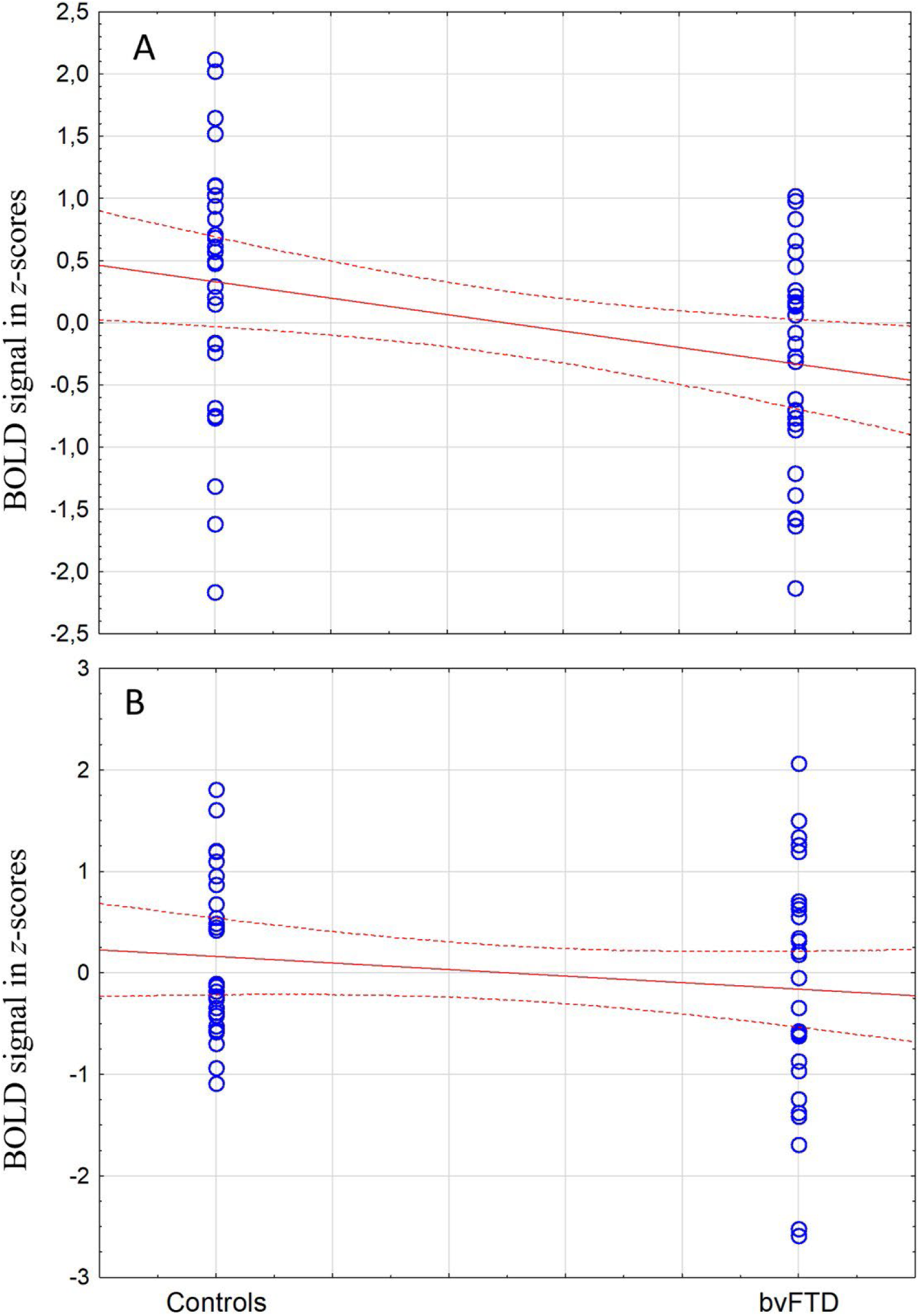
Significant difference between patients and controls in the right but not left insula in the affective perceptual empathy ROI. The figure displays difference between patients and controls in the right **(A)** and left **(B)** insula. Y-axis denote percent signal change during EFP converted to *z*-scores, x-axis diagnosis. Controls = controls subjects; bvFTD = the behavioral variant of frontotemporal dementia. Percent signal change in the right (*p =* 0.01, but not left *p=*0.23) is significant decreased in patients compared to controls.

**Table S1.** Clusters with significant reduced cortical thickness in patients compared with controls.

| lh | Size | X | Y | Z | CWP | Annot |
| --- | --- | --- | --- | --- | --- | --- |
|  | 42999.48 | -20.2 | 28.9 | 42.3 | 0.00010 | superiorfrontal |
|  | 3402.25 | -17.8 | -56.0 | 16.4 | 0.00010 | precuneus |
| rh | 13721.49 | 44.2 | -4.8 | -28.4 | 0.00010 | inferiortemporal |
|  | 4044.41 | 48.4 | -47.9 | 41.6 | 0.00010 | inferiorparietal |
|  | 2713.84 | 19.6 | 38.2 | 32.9 | 0.00010 | superiorfrontal |
|  | 1870.34 | 35.7 | 4.8 | 31.0 | 0.00010 | caudalmiddlefrontal |
|  | 1488.87 | 7.1 | -5.0 | 52.5 | 0.00010 | superiorfrontal |
|  | 489.42 | 45.1 | 28.3 | 27.3 | 0.02240 | rostralmiddlefrontal |
|  | 441.34 | 22.1 | -52.3 | 61.3 | 0.04060 | superiorparietal |

lh = left hemisphere; rh = right hemisphere; Size = size of clusters in $\text{mm}^2$ ; x, y, z coordinates; *CWP* = clusterwise *p*-value; Annot = cortical regions according to <sup>7</sup>.

**Table S2.** Subcortical volume differences between control subjects and patients with the behavioral variant of frontotemporal dementia (bvFTD)

| | *p-value* | $\eta^2$ |
| --- | --- | --- |
| Left_Lateral_Ventricle | p<0.01 | 0.15 |
| Left_Thalamus_Proper | p<0.01 | 0.13 |
| Left_Caudate | p<0.01 | 0.23 |
| Left_Putamen | p<0.01 | 0.36 |
| Left_Pallidum | p=0.04 | 0.08 |
| v3rd_Ventricle | p<0.01 | 0.12 |
| Left_Hippocampus | p<0.01 | 0.33 |
| Left_Amygdala | p<0.01 | 0.41 |
| Left_Accumbens_area | p<0.01 | 0.31 |
| Right_Lateral_Ventricle | p=0.01 | 0.11 |
| Right_Thalamus_Proper | p=0.01 | 0.11 |
| Right_Caudate | p<0.01 | 0.23 |
| Right_Putamen | p<0.01 | 0.41 |
| Right_Hippocampus | p<0.01 | 0.30 |
| Right_Amygdala | p<0.01 | 0.41 |
| Right_Accumbens_area | p<0.01 | 0.41 |

The table denotes difference in subcortical volumes between patients and controls, Effect size ( $\eta^2$ = partial eta squared).

**Table S3.**
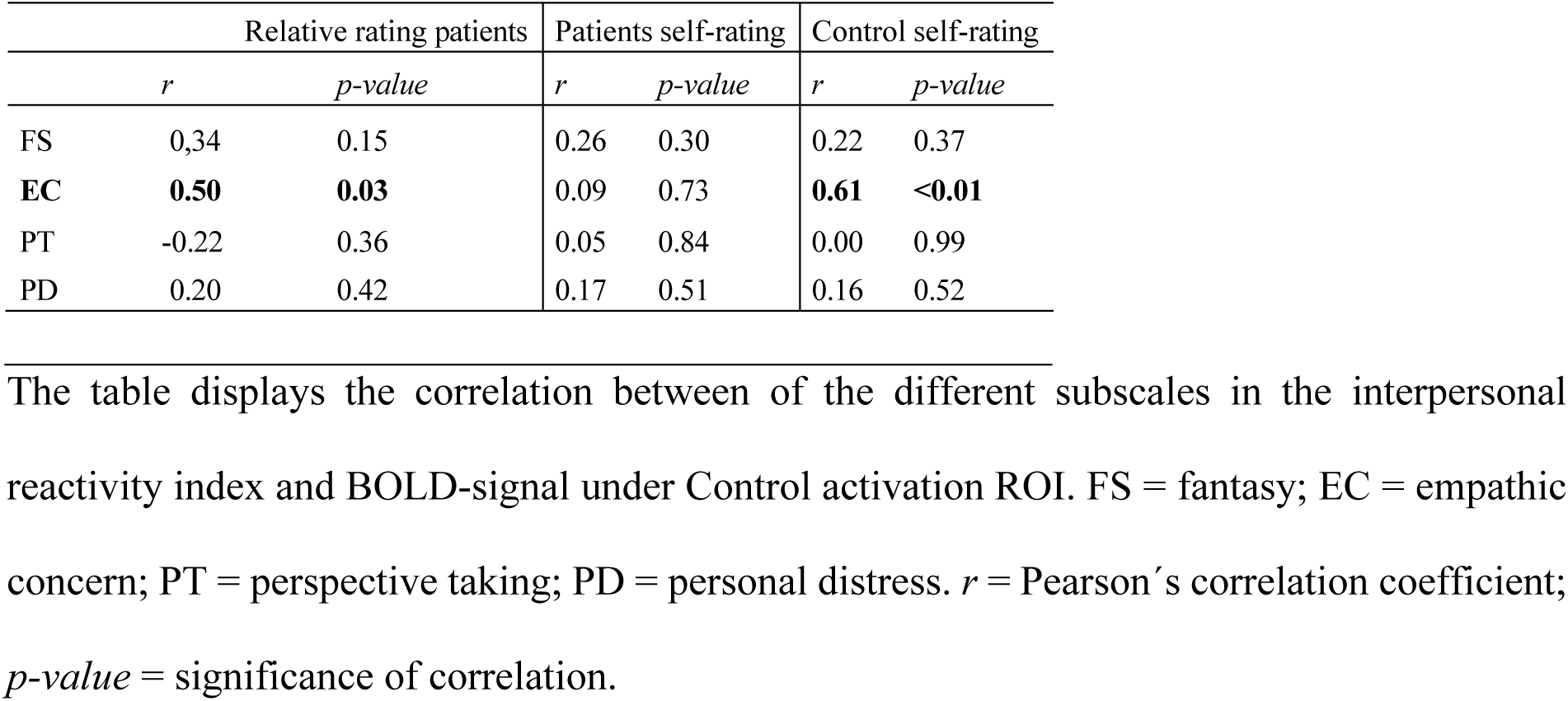
Correlation between mean BOLD-signal under the Control activation-ROI and the IRI sub-scales in patients and controls.

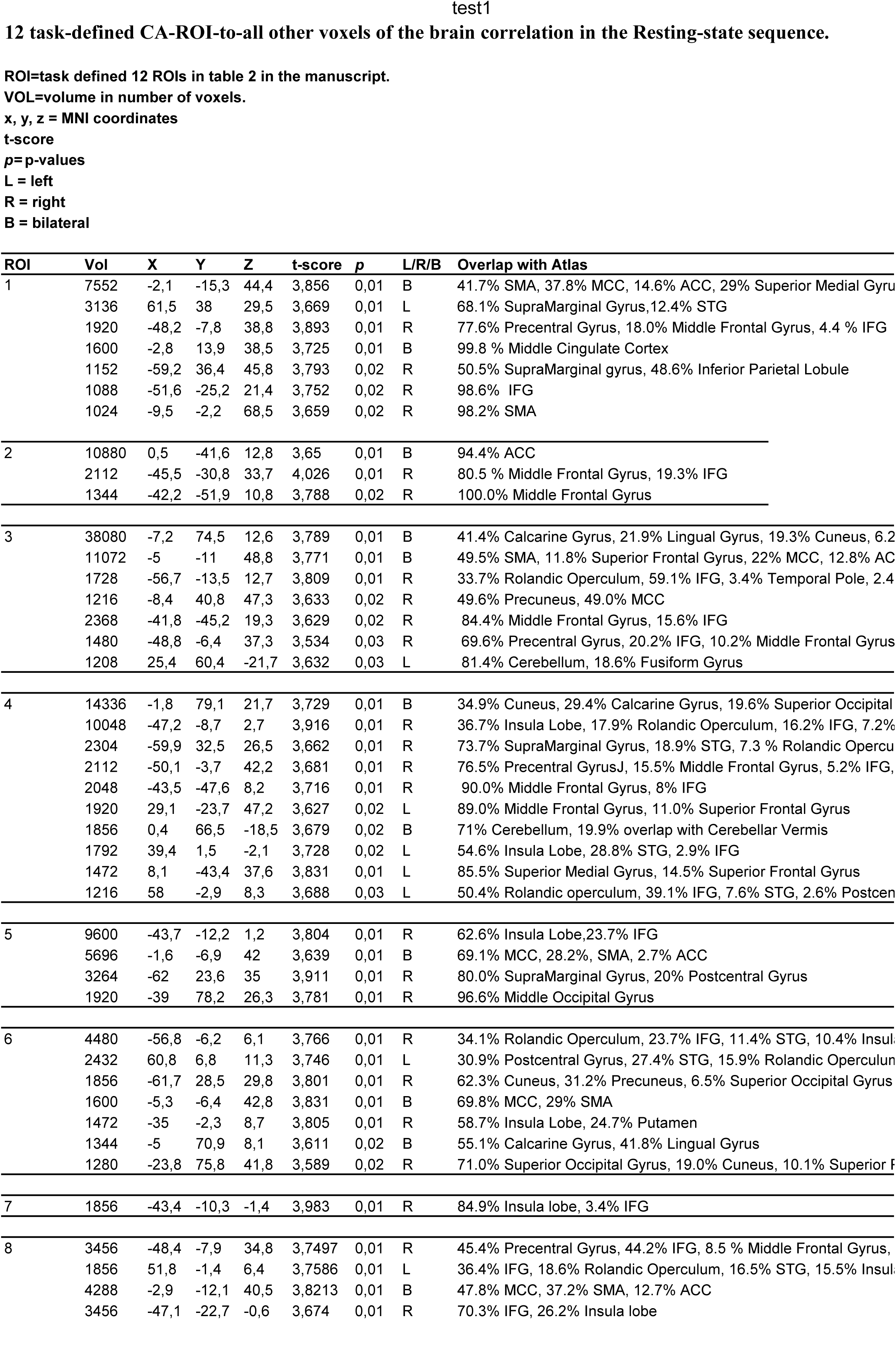
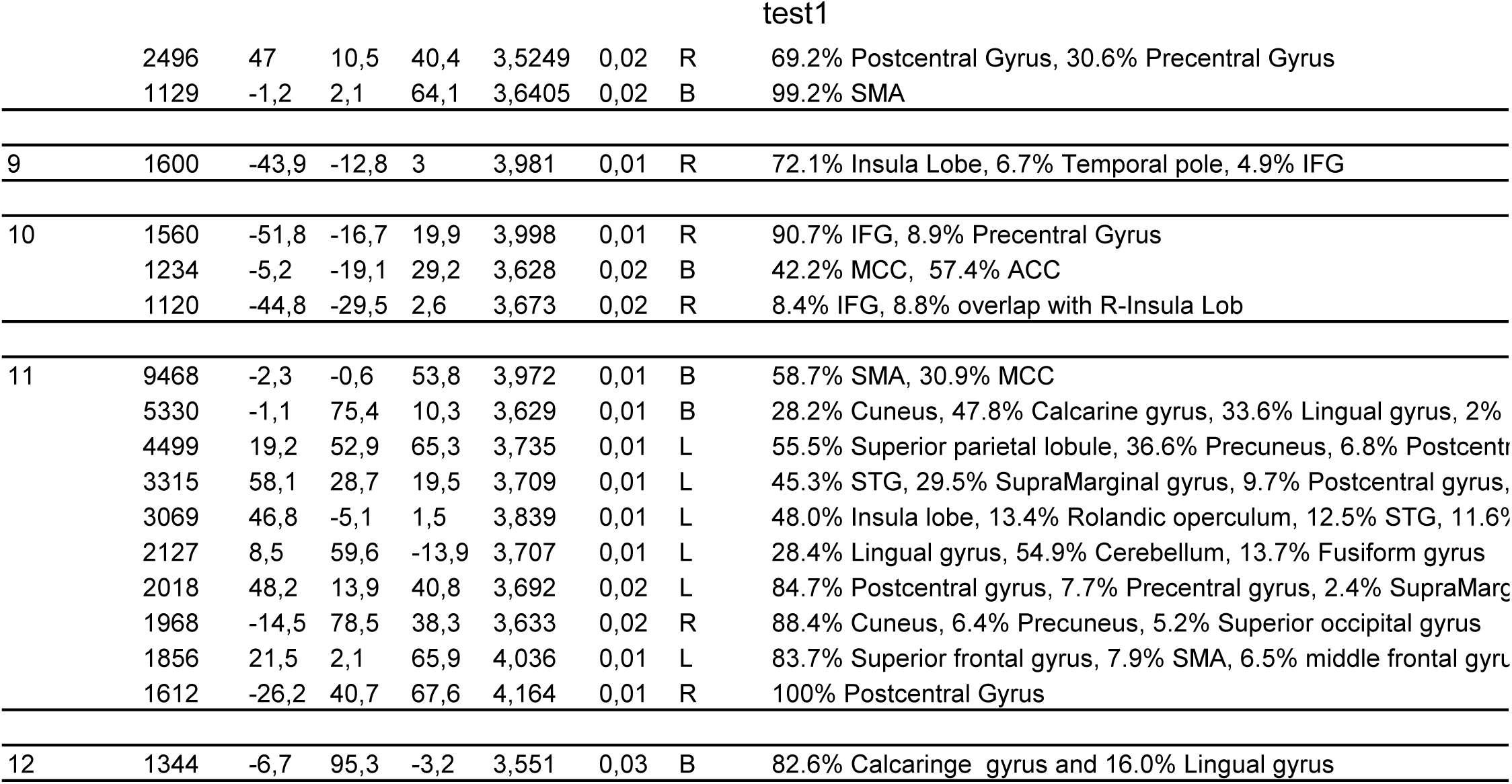

